# Cortical encoding of probabilistic temporal predictions during speech perception

**DOI:** 10.64898/2026.08.16.745095

**Authors:** Laure Deyna, Philippe Albouy, Agnès Trébuchon, Daniele Schön, Benjamin Morillon, Pierre Hieu Guilleminot

## Abstract

The temporal structure of speech has traditionally been characterized by the rhythmicity of its canonical linguistic units (phonemes, syllables, words), each summarized by a mean occurrence rate. While valid, this view overlooks whether speech carries a finer, context-dependent and probabilistic temporal structure that could support temporal predictive coding during listening. Using large French and English speech corpora, we trained models of increasing complexity to predict the onsets of linguistic units. Recurrent neural networks (RNNs) outperform mean-rate and hazard-rate models, showing that the variability around these rates is not noise but a temporal structure shaped by local context, statistically predictable across phonemes, syllables and words. Recording from 7,698 intracerebral electrodes in 53 neurosurgical patients listening to natural speech, we next show that the models’ output), the continuous probability of an upcoming onset (when), explains neural activity beyond acoustic and linguistic content (*what*) features, with markedly stronger effects for RNNs than for mean- or hazard-rate models. This dynamic neural prediction of *when* an onset will occur is dissociable from the encoding of linguistic content, relying on largely distinct channel populations. Temporal predictions engage a distributed cortical network extending from bilateral temporal cortex into left frontal and sensorimotor regions. Together, these results establish temporal prediction in speech as a dynamic, context-dependent and probabilistic process in its own right.

**Significant Statement:** When we listen to speech, the brain must anticipate not only *what* comes next but *when* it will occur. The what has been modeled within a dynamic long-term context; the when has mostly been characterized by a static temporal context through its periodicity. Analysing large speech corpora, we show that the onsets of linguistic units are predictable beyond their mean rate, carrying contextual, dynamic and probabilistic information. Using intracerebral recordings in patients listening to spoken stories, we demonstrate that the brain continuously encodes this temporal information, in neural populations largely separate from those that respond to linguistic content. These results place temporal structure at the heart of speech prediction, extending predictive coding from *what* is said to *when* it unfolds.

## Introduction

Speech perception is a fundamentally predictive process: as an auditory stream unfolds in real time, the brain must continuously anticipate both the *what* and the *when* of upcoming speech constituents. These two predictive dimensions have largely been studied as separate systems. The *what* has been captured by statistical models of linguistic identity, from early chain and n-gram models (1–3) through probabilistic surprisal models (4–6) to deep language models that specify each constituent as a function of its position within a rich, dynamically evolving context (7–10). The *when*, by contrast, has mainly been characterized through a quasi-regular rhythmic structure of speech and its neural tracking (11–16), an account that operates over the unfolding signal as a whole, largely independent of local temporal context. Here, we investigate purely temporal predictions during natural speech processing and aim to bridge the conceptual and methodological gap between the *what* and *when* frameworks. We extend accounts of the *when* beyond a global periodic structure to incorporate a local temporal context, driven purely by the timing of preceding constituents and never by their content.

Temporal expectations are known to modulate sensory gain, facilitate detection, and guide selective attention (17–21), and they are thought to be especially relevant for hierarchically organized auditory streams such as music and speech (22–25). Although speech is not perfectly isochronous, it exhibits robust quasi-regular rhythms and temporal prediction has been primarily framed through the preferred timescales at which linguistic units unfold: phonemes around 10-15 Hz, syllables around 4–5 Hz, content words around 1–2 Hz, or slower supra-lexical structures such as phrases, sentences, and intonational units (11, 12, 15, 16, 26). This framing is well-motivated from a neural perspective: the speech envelope shows a robust spectral peak between 2 and 8 Hz, closely aligned with syllabic rate (27, 28), and neural activity in auditory cortex reliably tracks this rhythm (29, 30), in a way that correlates with intelligibility (14). A parallel line of work has identified the dorsal auditory-motor pathway as a source of delta- and beta-band activity that supplies temporal predictions to auditory regions (31–34). Across these accounts, temporal prediction in speech is operationalized as alignment to the mean rate of a linguistic level (12, 26, 35).

Yet beyond these regularities, inter-onset intervals vary substantially around any central tendency and whether a systematic structure underlies this variability is still unclear (36–39). Outside speech, the brain is known to form temporal predictions from the statistical structure of preceding events even in the absence of strict periodicity: Such contextual predictions reflect the full distribution of expected event timing (40–42), are continuously updated by local context (43–46), and engage parietal and motor regions (47, 48). Critically, it remains unknown whether the temporal variability alone carries sufficient structure to support local, probabilistic temporal predictions in speech. Just as language models predict *what* word comes next with fine-grained probability, the question is whether a purely temporal model can predict *when* a word will occur. If so, further questions are which linguistic units are temporally predicted and whether the brain encodes such temporal probabilities in a way that is neurally dissociable from content predictions.

Here, we address these questions by combining large-scale speech corpora (French and English audiobooks) with human intracerebral recordings from 53 patients acquired during natural speech perception. To isolate temporal structure, we represented the presence of words, syllables and phonemes as binary onset timeseries, retaining timing alone, with no acoustic or lexical information. We trained a family of models of increasing contextual complexity, from probability density functions (PDFs) to hazard rates and recurrent neural networks operating on single (RNN-1D) or joint (RNN-3D) onset streams across phonemes, syllables, and words. We characterized, at each timestep, the probability of a unit onset given the past temporal context. We then used the model-derived probabilities as regressors of cortical activity recorded while participants listened to an audiobook. We show that linguistic timing is highly predictable and robustly encoded in bilateral temporal cortex, extending into left frontal and sensorimotor regions, in neural populations partially dissociable from those encoding speech content, establishing probabilistic timing as a distinct dimension of cortical speech processing.

## Results

### RNN models best capture temporal regularities

To assess whether onsets of linguistic units in natural speech can be predicted despite their temporal variability, we first extracted phoneme, syllable, and word onsets from a 25-hour French audiobook corpus (Fig. 1a). Onset distributions were unimodal and leptokurtic, with inter-onset intervals (IOIs) becoming longer and more variable from phonemes (median 80 ms; SD 150 ms) to syllables (180 ms; 234 ms) to words (260 ms; 323 ms), indicating a preferred timescale for each linguistic unit, with substantial variability around it. We then compared four models in their ability to predict the timing of the next linguistic onset: two standard approaches (probability density function, PDF, and hazard rate, HR) (41, 42, 49) and two recurrent neural networks (RNN-1D and RNN-3D; Fig. 1b,c). All four models take the past temporal context as input, encoded as the onsets of linguistic units, and return the probability of an onset occurring at each time step (fs = 50 Hz). The PDF, HR, and RNN-1D models each operate on a single linguistic unit at a time (one dimension). The PDF model captures the distribution of inter-onset intervals; the HR model extends this approach by incorporating the elapsed time since the last onset; and the RNN-1D model captures more complex temporal patterns through its recurrent units. The RNN-3D model is identical to the RNN-1D but simultaneously takes the temporal information of all three linguistic units (phonemes, syllables, words) as input, and thus benefits from interactions between their temporal structures. This allowed us to test whether onset timing is predictable at all, and to identify which hypotheses, ranging from mean rate (PDF) to contextual dependencies across hierarchical levels (RNN-3D), best account for the observed variability.

**Figure 1.**
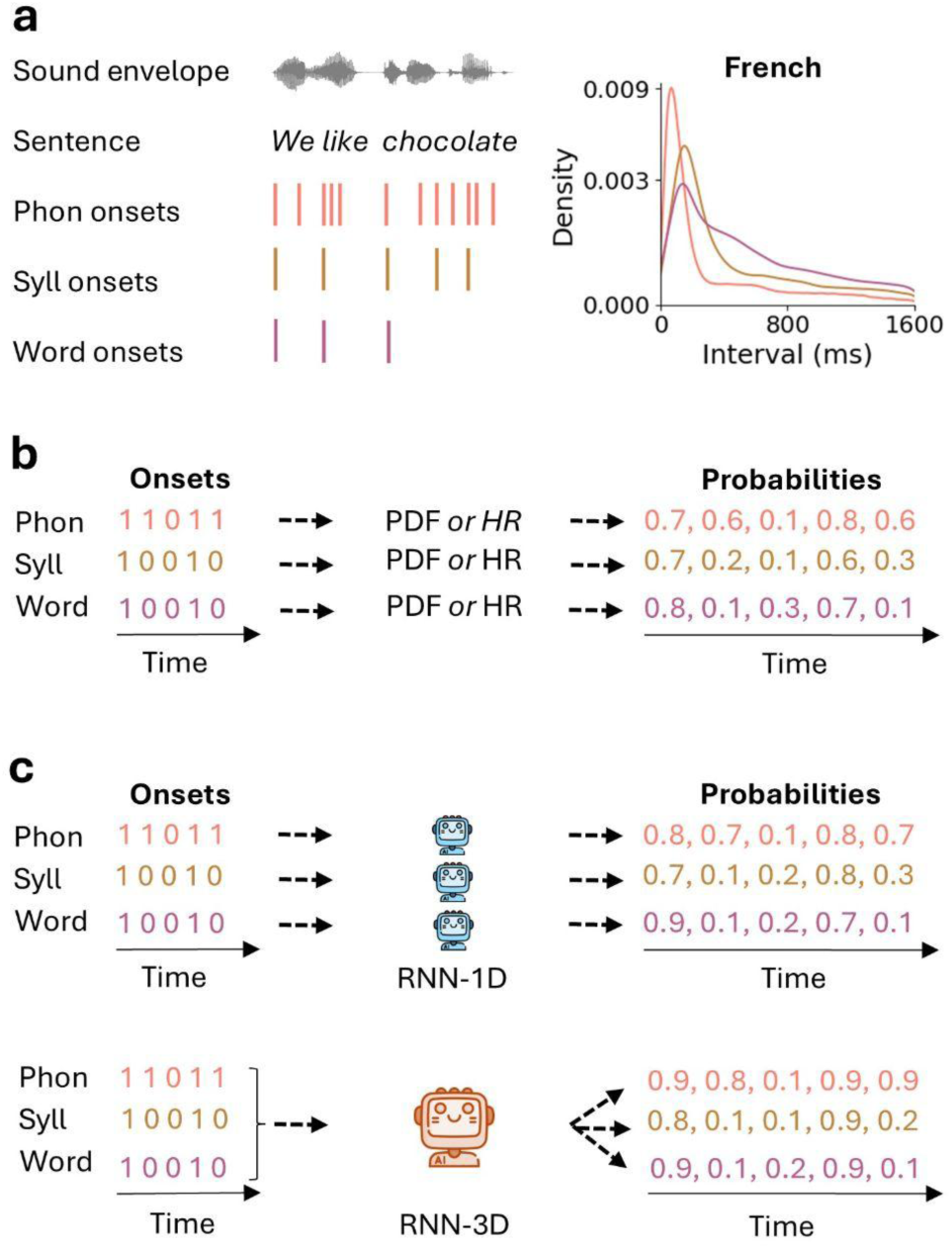
Construction of Inter-Onset Interval (IOI) speech corpora and modeling of onset prediction. **(a)** Left: Example of onset extraction for linguistic units (phonemes, syllables, and words) from a spoken sentence. Right: Distributions of IOIs in the French training corpus (25 h). The probability density (y-axis) is displayed on a semi-logarithmic scale; the time (x-axis) is truncated at 1600 ms. **(b–c)** Input and output structures of the models used to temporally predict linguistic onsets. **(b)** Standard models: Each linguistic unit (phoneme, syllable, word) is processed independently. The input is a binary time series indicating the presence (1) or absence (0) of an onset for a given unit, up to the current time step. The output is the predicted probability of an onset occurring at the next time step (sampling frequency, fs = 50 Hz). The model can be either a probability density function (PDF), describing the distribution of onsets over time, or a hazard rate (HR), the instantaneous probability of an onset given it has not yet occurred (reset after each occurrence). **(c)** RNN models: RNN-1D is a one-dimensional recurrent neural network, trained and tested separately for each linguistic unit. In contrast, RNN-3D jointly processes all three linguistic unit time series as a 3-dimensional input and outputs the corresponding onset probabilities for each unit.

All models significantly predicted the timing of upcoming onsets above chance (AUC chance level = 0.5; Wilcoxon signed-rank test with Bonferroni correction, all p < 0.001, Rank Biserial Correlation (RBC) ≃ 1.0; Fig. 2a), which shows that the temporal structure of speech is predictable from onset history alone. A Friedman test revealed a robust main effect of model type for each linguistic unit (all χ²(3) > 291, all p-unc < 0.001). Post-hoc comparisons showed a graded hierarchy of performance that follows model complexity, fully consistent across linguistic units (for all pairwise model comparisons, p < 0.001, Nemenyi corrected, Fig. 2a). The PDF model performed well above chance, and the HR model performed significantly better.. The RNN-1D model outperformed these standard models, demonstrating the importance of prior temporal context beyond rate information (PDF) and previous onset (HR) in predicting the next onset. Finally, the RNN-3D model, that jointly processes phoneme, syllable, and word onset sequences, achieved the highest performance (up to AUC = 0.83 for one syllabic speech segment), highlighting that the temporal structure of speech relies on interactions across linguistic levels.

**Figure 2.**
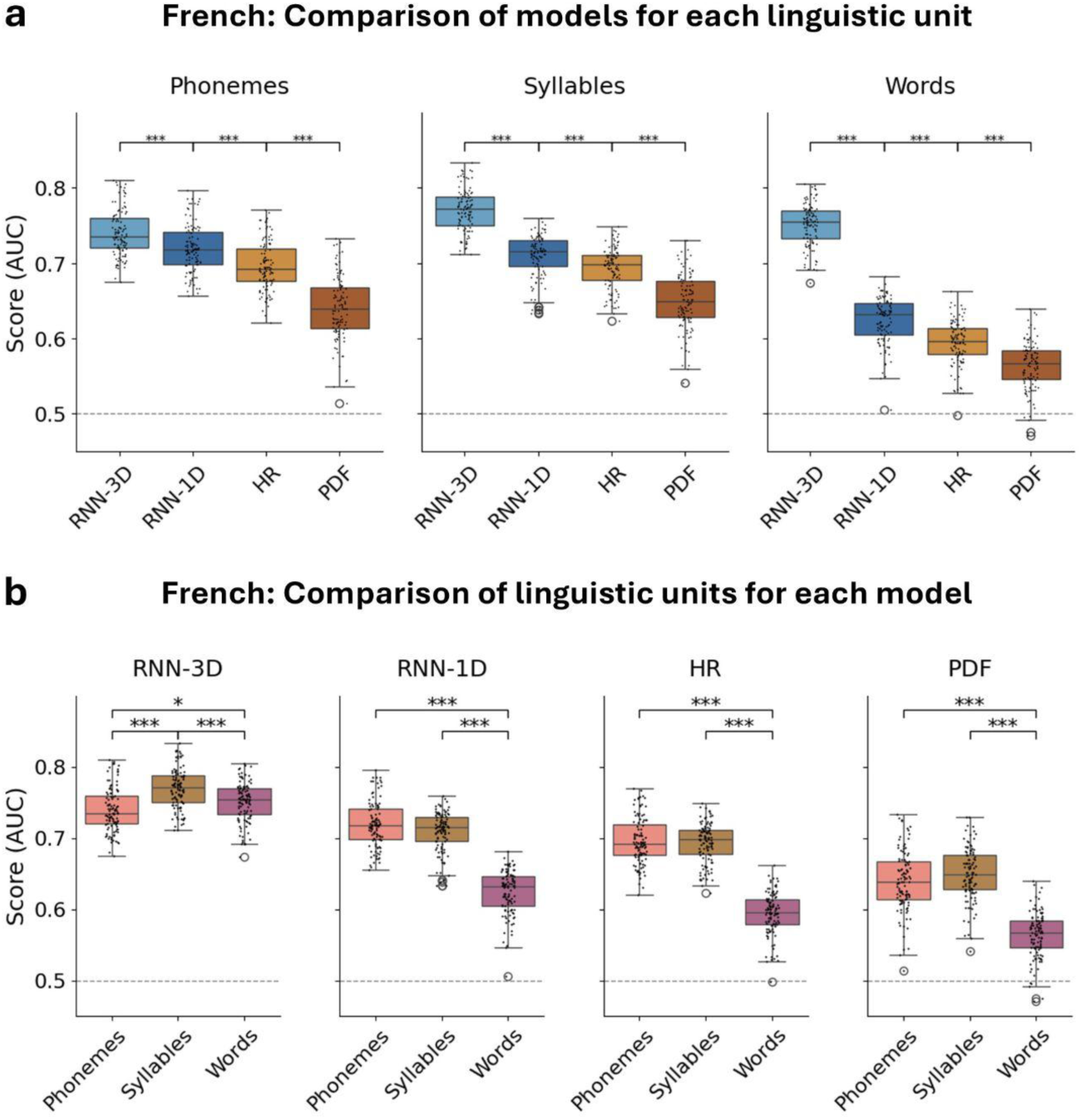
Comparison of model performance across linguistic units in French. Distribution of area under the ROC curve (AUC) scores computed on 100 one-minute speech segments. Models were evaluated based on their ability to temporally predict the onset of upcoming linguistic units (phonemes, syllables, or words). **(a)** AUC score comparison between models, for each linguistic unit. **(b)** AUC score comparison between linguistic units, for each model. Black dots represent individual speech segments (n = 100; *p < 0.05, ***p < 0.001, Nemenyi post-hoc tests). Boxplots show the median, interquartile range (IQR; box) and whiskers at 1.5× IQR. The horizontal dashed lines indicate chance level (0.5). Model labels: RNN-3D (RNN trained on all linguistic units), RNN-1D (RNN trained separately for each linguistic unit), HR (hazard rate), PDF (probability density function).

### Cross-unit interactions enhance temporal predictability

We next compared the temporal predictability of phoneme, syllable, and word onsets (Fig. 2b). A Friedman test revealed a significant effect of linguistic unit for each model (all χ²(2) > 132, p-unc < 0.001), and post-hoc Nemenyi comparisons showed a consistent hierarchy across models. For the models predicting each unit independently (PDF, HR, and RNN-1D), phonemes and syllables were predicted with comparable accuracy (all p > 0.1, Nemenyi corrected), and both were significantly more predictable than words (all p < 0.001, Nemenyi corrected), likely reflecting the broader inter-onset interval distribution of words (see above, Fig. 1a).

In contrast, the RNN-3D model (Fig. 2b) revealed a distinct profile. Syllables were significantly more predictable than both phonemes and words (all p < 0.001, Nemenyi corrected), while words were more predictable than phonemes (p < 0.05, Nemenyi corrected). This suggests that syllables, as intermediate units, benefit most from cross-level interactions, integrating information from both finer (phoneme) and coarser (word) timescales. The improved predictability of words compared to phonemes further suggests that higher-level units, despite greater intrinsic variability (see Fig. 1a), can exploit hierarchical temporal context.

### Temporal predictability replicates in English

To assess whether these findings are robust across languages, we trained and evaluated architecturally identical models on a 25-hour English audiobook corpus. IOI distributions were comparable to those in French: shorter and more regular for phonemes (median = 80 ms, SD = 98 ms), intermediate for syllables (median = 180 ms, SD = 181 ms), and longer and more variable for words (median = 260 ms, SD = 252 ms). All models predicted phoneme, syllable, and word onsets significantly above chance (Wilcoxon signed-rank with Bonferroni correction, all p < 0.001, RBC ≃ 1.0; Fig. S1). Comparisons across models revealed the same hierarchy as in French (Friedman tests, all χ²(3) > 285, all p-unc < 0.001, and post-hoc analyses all pairwise p < 0.001, Nemenyi corrected; Fig. S1a). Analyses across linguistic units further showed significant effects for each model (Friedman tests, all χ²(2) > 147, all p-unc < 0.001; Fig. S1b). However, the pattern for the RNN-3D model diverged between languages: in English, words were the most predictable units (p < 0.001, Nemenyi corrected), whereas syllables dominated in French (see Fig. 2b). These findings demonstrate that for both French and English, the onsets of linguistic units are best predicted by models exploiting the complex temporal structure of speech, although the unit benefiting most from hierarchical context seems language-dependent.

### The brain encodes temporal predictions during naturalistic speech listening

Having established that the timing of linguistic-unit onsets in natural speech is predictable from past temporal context, we next asked whether the brain encodes this temporal predictability during speech perception. We analysed intracerebral recordings (stereotactic EEG) obtained while patients (n = 53; total channels = 7,698) listened to a 10-minute French audiobook, using an encoding approach (temporal response function, TRF; (50)). Bipolar signals were filtered in the low-frequency range (0.3–24 Hz), which best captures the cortical tracking of speech dynamics (51–55) as well as the anticipatory activity supporting temporal prediction (33, 41, 42). We used a unique variance scheme to test whether temporal predictability was encoded beyond the acoustic and linguistic features typically involved in speech processing (51, 56). As control features, we included binary onset time series, acoustic features capturing low-level energy dynamics and acoustic proxies of phonemic-to-phrasal linguistic timescales, and linguistic content features, namely the surprisal and entropy associated with words and phonemes (Methods; Fig. 3a, S2 and S3).

**Figure 3.**
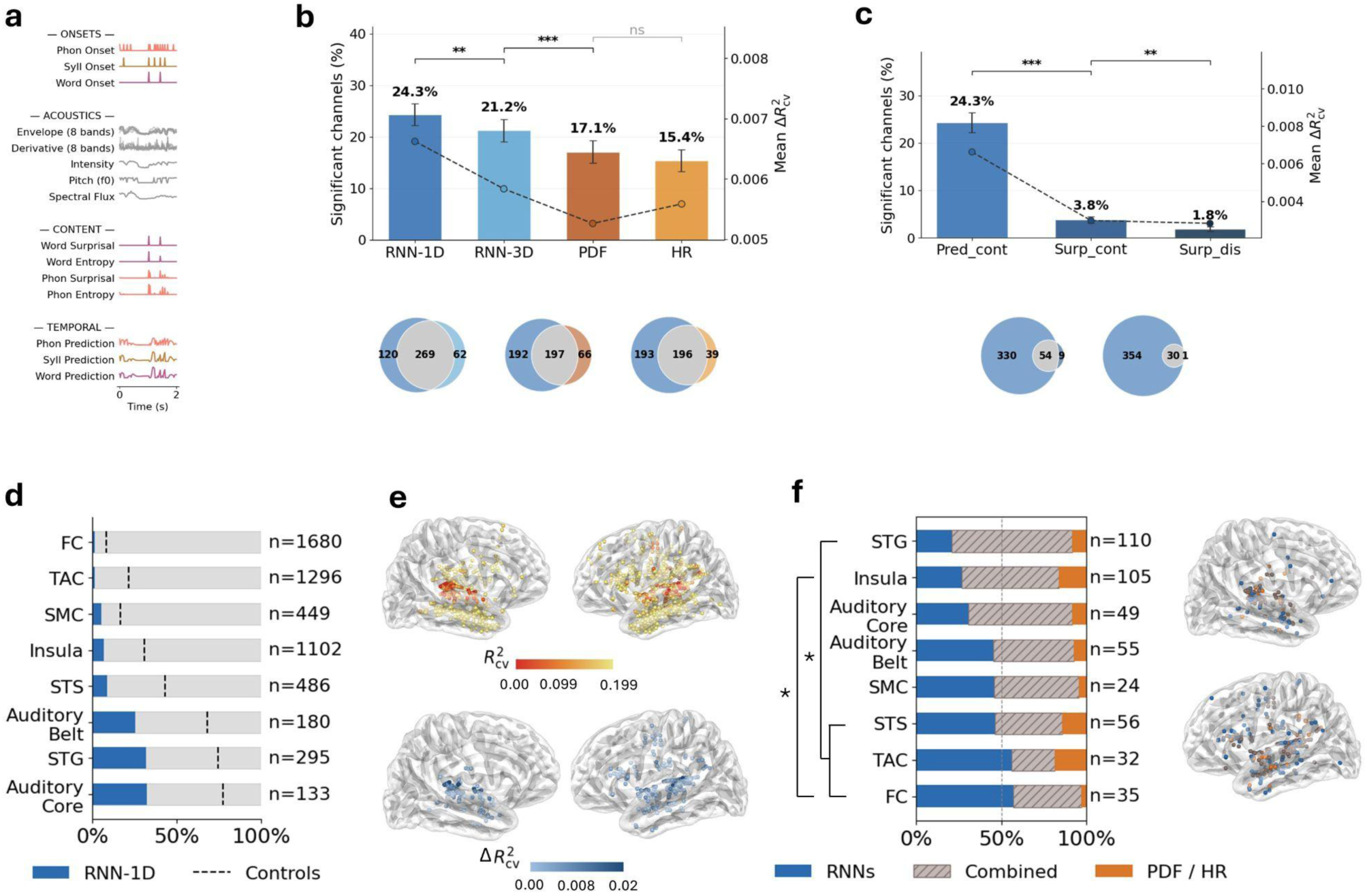
Neural encoding of temporal predictions during speech perception. **(a)** Features entered as regressors in the temporal response function (TRF) analysis, shown over a 2 s excerpt. Onsets: binary onset time series for phonemes, syllables and words. Acoustics: an 8-band spectrogram envelope and its 8-band derivative (acoustic edges), intensity, pitch strength (f0) and spectral flux. Content: surprisal and entropy for words and phonemes. Temporal: model-derived onset predictions for phonemes, syllables and words (temporal-prediction regressors). **(b)** For each model, percentage of speech-responsive channels (significant for the control model: onsets, acoustics and content features; Table 1) where its temporal-prediction regressors explained significant unique variance beyond it. Each model’s phoneme, syllable and word onset predictions are entered jointly as three regressors. Bars show the mean ± SEM across participants; the dashed line (right axis) shows the mean unique explained variance (ΔR²_cv). Models: RNN-1D (dark blue), RNN-3D (light blue), PDF (rust), HR (orange). Across-model comparisons: pairwise Wilcoxon tests on adjacent models, Bonferroni-corrected. Venn diagrams: overlap in significant channels between RNN-1D and each of the other three models. Channels pooled across participants. **(c)** Same as (b), for three metrics derived from RNN-1D: the continuous onset-prediction signal (Pred_cont, used in b) and two surprisal-based metrics (Surp_cont, continuous surprisal; Surp_dis, discrete surprisal). Venn diagrams: overlap in significant channels between Pred_cont and each of the two surprisal metrics. **(d)** For each region, proportion of channels significant for the RNN-1D model (blue bar) out of all channels in the region (pale bar; n per region). Dashed vertical lines: proportions of speech-responsive channels. Regions are ordered by increasing RNN-1D proportion. **(e) Top.** Spatial distribution of explained variance (R²_cv) of the speech-responsive channels; non-significant channels are not shown. **Bottom.** Channels significant for the temporal-prediction regressors (RNN-1D); color encodes ΔR²_cv. Colormaps scaled from 0 to the 95th percentile. **(f) Left.** Regional breakdown of channels with significant unique variance, by model family. Within each region, the proportion of significant channels that are RNNs only (RNN-1D or RNN-3D; blue), both families (Combined, hatched) or PDF/HR only (orange); n significant channels per region. Regions ordered by increasing RNNs-only proportion. Brackets and asterisks: between-region differences in the RNNs-only proportion (Fisher’s exact test, RNNs-only vs. Combined + PDF/HR, Bonferroni-corrected across all region pairs). **Right.** Spatial distribution of the channels shown on the left. **General conventions.** N = 53 participants (7,698 channels). Channel-level significance: FWER-corrected permutation test (max statistic, n = 1000, p ≤ 0.05); for unique variance (ΔR²_cv), only the added regressors were permuted. *p < 0.05, **p < 0.01, ***p < 0.001; ns, not significant. Model labels as in Figure 2. Region abbreviations (d, f): STG (superior temporal gyrus), STS (superior temporal sulcus), TAC (temporal associative cortex), SMC (sensorimotor cortex), FC (frontal cortex).

**Table 1.** Encoding models: neural encoding of temporal predictions (Fig. 3). *Acoustics = 8 spectrogram bands + 8 derivatives + spectral flux + intensity + pitch strength Onsets = word onsets + syllable onsets + phoneme onsets*

| Model | Regressors |
| --- | --- |
| Control | Acoustics + Onsets + word surprisal + word entropy + phoneme surprisal + phoneme entropy |
| RNN-1D | Control + RNN-1D predictions (word, syllable, phoneme) |
| RNN-3D | Control + RNN-3D predictions (word, syllable, phoneme) |
| PDF | Control + PDF predictions (word, syllable, phoneme) |
| HR | Control + HR predictions (word, syllable, phoneme) |
| Pred_cont | Control + RNN-1D predictions (word, syllable, phoneme) |
| Surp_cont | Control + RNN-1D continuous surprisal (word, syllable, phoneme) |
| Surp_dis | Control + RNN-1D discrete surprisal (word, syllable, phoneme) |

*Each model adds its regressor(s) on top of the control, which is held fixed (not permuted). $\Delta R^2_{cv} = R^2_{cv}(\text{model}) - R^2_{cv}(\text{control})$ . To obtain the speech-responsive channels (significant for the control itself), the entire control is permuted.*

Across participants, the TRF model computed over all these control features (Fig. 3a, Table 1) significantly explained neural responses in 22.8% of recording sites (hereafter speech-responsive channels, 1,752 channels; FWER-corrected permutation test, max statistic, p ≤ 0.05; mean R²_cv = 0.048, SD = 0.073, max = 0.615; Fig. 3d,e; regions defined in Fig. S4). The speech-responsive channels with the highest predictive accuracy were predominantly localized to the superior temporal gyrus (STG) bilaterally and, more broadly, to auditory and perisylvian regions, consistent with the established role of these areas in encoding acoustic and linguistic features of continuous speech (57, 58).

We then asked which model of onset timing best explained neural activity beyond control features. We compared the four models introduced above (PDF, HR, RNN-1D and RNN-3D), by adding, in separate analyses, each model’s temporal predictions to the control features. These temporal predictors consisted of the output of the models: continuous (i.e., at each timestep) probabilities of onset occurrence for phonemes, syllables and words (Fig. S5). We then identified the channels whose neural activity reconstruction was significantly improved by the addition of temporal-prediction features (FWER-corrected permutation test, max statistic, n = 1000, p ≤ 0.05). These significant channels were a subset of the speech-responsive channels (Fig. 3e and S6), so we expressed their number per participant relative to this set (Fig. 3b). The four models differed in the fraction of channels they rendered significant (Friedman χ²(3) = 61.5, p < 0.001). RNN-1D had the largest fraction of channels significant (24.3% of speech-responsive channels; 389 channels), followed by RNN-3D (21.2%; 331 channels), PDF (17.1%; 263 channels) and HR (15.4%; 235 channels). Pairwise comparisons on the number of significant channels (Wilcoxon signed-rank on adjacent models, Bonferroni-corrected) confirmed an advantage of RNN-1D over RNN-3D (p = 0.008) and of RNN-3D over PDF (p < 0.001), whereas PDF and HR did not differ (p = 0.07). The mean unique variance beyond the control model was likewise highest for RNN-1D (ΔR²_cv = 0.0066; Fig. 3b, right axis). The RNN-1D-significant channels encompassed most of those of the other models (269 shared with RNN-3D, 197 with PDF, 196 with HR; Fig. 3b, Venn diagrams). Notably, RNN-1D was the best model for neural regression despite being the simpler recurrent model and a less accurate onset predictor than RNN-3D at the speech corpus level (Fig. 2). When phoneme, syllable and word predictions were evaluated separately, the advantage of RNN-1D over RNN-3D held for syllables only (Fig. S7), and the unique contribution of each unit was small and comparable across units (Fig. S8). Corpus-level predictive accuracy therefore does not directly translate into relevance for neural encoding.

We next asked whether the brain encodes temporal predictions as raw probabilities, or instead as a form of surprisal, emphasizing prediction violations. We compared the continuous prediction with two surprisal-based metrics computed from the same output: continuous surprisal and discrete surprisal at linguistic-unit onsets (Methods), with the three linguistic units’ regressors entered jointly, as in the model comparison above. The untransformed continuous prediction rendered far more channels significant than either surprisal metric (24.3% vs. 3.8% and 1.8% of speech-responsive channels; Friedman χ²(2) = 87.9, p < 0.001; adjacent pairwise Wilcoxon signed-rank, Bonferroni-corrected: continuous prediction > continuous surprisal, p < 0.001; continuous surprisal > discrete surprisal, p = 0.003; Fig. 3c and S9). Accordingly, it also yielded the largest mean unique variance (ΔR²_cv = 0.0066, vs 0.0030 and 0.0028). We therefore used the untransformed continuous prediction as the temporal predictor throughout.

Finally, we asked whether contextually rich (RNN-1D, RNN-3D) and contextually poor (PDF, HR) models explain neural variance in the same brain regions. Grouping channels by model family (RNNs = RNN-1D ∪ RNN-3D, n = 451; PDF/HR = PDF ∪ HR, n = 327; Both = RNNs ∩ PDF/HR, n = 268 channels significant for both), we found that the two families were not encoded in identical areas (Fig. 3f). The proportion of channels significant for RNNs only, varied across regions (Fisher’s exact tests, Bonferroni-corrected): it was lowest in the STG, where significant channels were predominantly shared between families, and significantly higher in higher-order regions, including the superior temporal sulcus (STS; p = 0.03), the temporal associative cortex (TAC; p = 0.007) and the frontal cortex (FC; p = 0.003); the insula likewise contained fewer RNNs-only channels than FC (p = 0.050). Thus, while both families of models converged in the primary auditory/STG cortex, the contextually rich RNN models were comparatively more prominent than the context-poor PDF/HR models in higher-level cortical areas.

### Temporal prediction is dissociable from linguistic content prediction

The previous analyses demonstrated that during speech comprehension, the brain represents temporal predictions (*when* will a unit occur), beyond content prediction (*what* unit will occur). We then asked directly whether these two forms of prediction operate within distinct neural populations. Using the same TRF framework, we isolated the unique contribution of temporal prediction features (RNN-1D continuous predictions) and linguistic content prediction features (word and phoneme surprisal and entropy, (4, 9, 56)), using a symmetric variance-partitioning scheme in which each set served as a control for the other, in addition to the acoustic and onset regressors (Methods; Table 2). Speech-responsive channels were accordingly redefined as channels significant for the acoustic and onset features alone. Note that the two sets of predictive features were computed with different metrics: continuous onset probabilities for temporal prediction (Fig. 3a), and surprisal and entropy time-locked onto the relevant linguistic-unit onsets for content prediction. This distinction is intrinsic to each model: the RNN operates over time steps and natively outputs continuous onset probabilities, whereas LLMs operate over discrete tokens.

**Table 2.** Encoding models: temporal vs. content predictions (Fig. 4; word and phoneme levels only). *Acoustics = 8 spectrogram bands + 8 derivatives + spectral flux + intensity + pitch strength Onsets = word onsets + syllable onsets + phoneme onsets*

| Model | Regressors |
| --- | --- |
| *Control_for_temporal* | Acoustics + Onsets + word surprisal + word entropy + phoneme surprisal + phoneme entropy (*Control in Table 1*) |
| *Control_for_content* | Acoustics + Onsets + RNN-1D predictions (word, phoneme) |
| *Temporal* | *Control_for_temporal* + RNN-1D predictions (word, phoneme) |
| *Content* | *Control_for_content* + word surprisal + word entropy + phoneme surprisal + phoneme entropy |
| *Combined* | Acoustics + Onsets + RNN-1D predictions (word, phoneme) + word surprisal + word entropy + phoneme surprisal + phoneme entropy |

*Each effect uses the same full regressor set; only its control differs, defining its unique variance ( $\Delta R^2_{cv}$ ): the other predictor set for Temporal and Content, Acoustics + Onsets alone for Content+Temporal.*

Both forms of prediction uniquely contributed to the neural data (Fig. 4a,b,d). Channels displaying unique temporal contribution were more numerous than those displaying unique content contribution (22.4% vs 16.9% of control-significant channels, i.e., channels significant for the acoustic and onset features alone;; Wilcoxon signed-rank, p = 0.019). A joint model containing both predictor sets rendered more channels significant than either unique contribution alone (31.0%; Fig. 4a), consistent with variance shared between the two sets of predictors (Fig. S2). At the single-channel level, plotting each significant channel’s unique temporal gain against its unique content gain (Fig. 4c,d) revealed three largely separate populations: channels driven by temporal prediction only (n = 229), by content prediction only (n = 123), and by both unique variances (n = 122), with the unique gains falling close to one or the other axis rather than co-varying.

**Figure 4.**
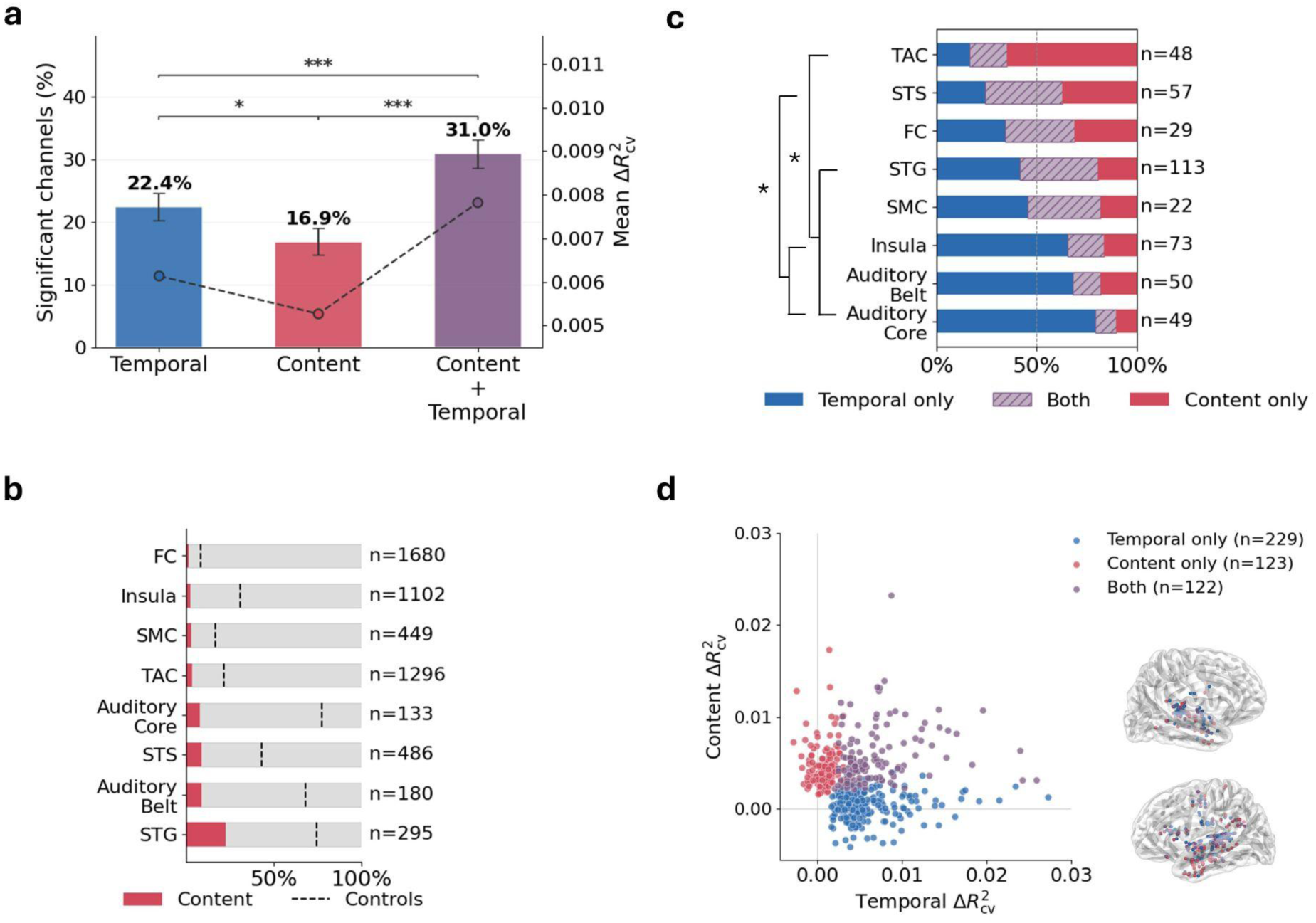
Neural encoding of temporal vs content predictions. **(a)** For control-significant channels (significant for the onset and acoustic features; content excluded, unlike the speech-responsive channels of Figure 3), percentage with significant temporal unique variance (blue), content unique variance (red), or variance from a single TRF trained on both predictor sets (Content+Temporal; purple). The three overlap, so a channel may enter more than one bar. Temporal and content unique variances are estimated over a control that also includes the other predictor set (Methods). The Content+Temporal model can be significant in channels where neither unique variance is (omitted from (c) and (d)). Across-condition comparisons: pairwise Wilcoxon signed-rank tests, Bonferroni-corrected. Dashed line (right axis): mean ΔR²_cv. **(b)** For each region, proportion of channels with significant content unique variance (red bar) out of all channels in the region (pale bar; n per region). Dashed vertical line: proportion of control-significant channels. Regions ordered by increasing content proportion. **(c)** Channels with significant temporal and/or content unique variance, split into three mutually exclusive categories: temporal only (blue), content only (red) or both (purple, hatched). Within each region, proportion of significant channels in each category (n per region). Regions ordered by increasing temporal-only proportion. Brackets and asterisks: Fisher’s exact test comparing temporal-only vs content-only counts (Bonferroni-corrected across all region pairs). **(d) Left.** Per-channel temporal and content unique variance (ΔR²_cv): temporal (x-axis) vs content (y-axis). Categories and colors as in (c): temporal only (blue, n = 229), content only (red, n = 123), both (purple, n = 122). **Right.** Spatial distribution of these channels on the reconstructed cortical surface. **General conventions.** Temporal predictions here comprise the RNN-1D phoneme and word predictions (syllable excluded, to match the content metrics; Methods). Conventions as in Figure 3.

This functional dissociation had a clear anatomical counterpart (Fig. 4c,d). The relative balance of temporal-only versus content-only channels differed across regions (Fisher’s exact tests, Bonferroni-corrected): temporal-prediction channels predominated in primary auditory cortex (Auditory Core and Belt) and the insula, whereas content-prediction channels were comparatively over-represented in higher-order temporal and temporal associative cortex. In particular, the temporal associative cortex (TAC) and the superior temporal sulcus (STS) contained significantly higher proportions of content-only channels than the auditory core, auditory belt and insula (TAC and STS vs these auditory regions, all p < 0.05; TAC vs STG, p < 0.001). This pattern describes a gradient from primary auditory regions, dominated by temporal prediction, to higher-order regions, where content prediction becomes prominent. Together, these results show that while the two types of prediction share part of their explained variance, they remain dissociable processes supported by partly distinct cortical networks.

## Discussion

Our study reveals the temporal structure of speech as a complex and predictable source of information, by showing that: (1) onsets of linguistic units are predictable beyond their mean occurrence rate; (2) this predictive process relies on local temporal context and inter-unit interactions, as captured by recurrent neural networks (RNNs); (3) the brain makes temporal predictions during natural speech listening, as the continuous probability of onset occurrence is encoded in neural activity beyond acoustic and content features; and (4) this temporal prediction is dissociable from linguistic content prediction, recruiting partly distinct cortical networks along the auditory processing hierarchy.

At the corpus level, the ability of the PDF and HR models to predict upcoming phoneme, syllable and word onsets confirms the existence of preferred timescales for each of these linguistic units, consistent with their previously hypothesized quasi-periodic structure (15, 16). However, the greater performance of the RNNs shows that the variation around these timescales is not noise but carries contextual, dynamic and probabilistic information. Indeed, RNNs were chosen for their ability to capture temporal patterns through recurrent population dynamics (59, 60), thus generating predictions that account for long-range temporal dependencies. Notably, RNN-3D was the best-performing model for every linguistic unit in both French and English, showing that the temporal structure of each unit also depends on its interactions with the others. It is not possible, however, to determine which type of temporal structure the RNNs capture: simple repeating patterns, more complex sequences or, for RNN-3D, cross-unit dependencies. What we can affirm is that (1) the purely temporal structure of speech carries predictable information, and (2) this information exists both within each linguistic unit and in the interactions between units. The linguistic units best predicted differ depending on the language and the model used, leaving open how language-specific temporal structure shapes predictability (13, 36, 38, 61).

At the neural level, our results show that the temporal information captured by the models’ predictions at the corpus level is encoded by the brain during speech listening. The predictions derived from the PDF and HR models were significantly encoded, in line with the established role of rhythm in sensory (20, 33) and, specifically, speech processing (12, 62). Crucially, the models rendering the largest number of significant channels were the RNNs: the brain encodes local contextual variations beyond temporal regularities. Although the RNN-3D model is unambiguously the best at predicting onsets at the corpus level, it is not the best model for explaining neural activity. This result should be interpreted with caution, however: the channels responsive to RNN-1D and RNN-3D overlap substantially, and when linguistic units are considered separately, RNN-1D outperformed RNN-3D only for syllables (Fig. S7). This pattern likely reflects the strong correlation between RNN-1D and RNN-3D predictions (Fig. S2), and may also relate to the observation that a model’s predictive accuracy does not systematically translate into its neural predictivity (63), but see (10). A related question is whether any linguistic unit dominates the neural encoding (Fig. S8). When each unit was considered in isolation, syllable predictions rendered the most channels significant, echoing the prominence of the syllabic timescale in auditory cortical tracking (12, 15). The units’ unique contributions, however, were small, statistically indistinguishable, and carried by largely non-overlapping channel populations. Temporal predictions thus appear to be largely shared across linguistic levels, with no unit uniquely dominating. This is expected if the brain tracks the predicted timing of each linguistic level, rather than privileging the most predictable one.

Importantly, these temporal predictions were best modelled as the continuous probability of onset occurrence rather than time-locked to unit onsets or logarithmically transformed into surprisal (Fig. 3c and S9). In this sense, temporal prediction appears to be a distinct mechanism from content prediction, which is time-locked to the unit’s onset and traditionally modeled via surprisal (4, 8, 9). Whereas surprisal is jointly determined by the prediction and the actual input, and thus reflects the outcome of the predictive process, the continuous probability characterizes the prediction itself, defined at every moment including between onsets. The brain thus appears to maintain a continuously updated estimate of when the next linguistic units will occur: a prediction signal proper, rather than only the prediction errors emphasized in predictive coding studies (64). This contrasts with the encoding of the acoustic signal itself, which is best captured by discrete, event-like representations: auditory cortical responses track acoustic edges, the brief moments of sharp energy rise that mark vowel and syllable onsets, rather than the continuous envelope (62, 65).

In light of our results as a whole, we hypothesize that temporal prediction constitutes a facilitation process, consistent with dynamic attending accounts of temporal structure (18, 66). Predictions derived from the simplest models (PDF, HR), based on the regularity of linguistic units, were mostly encoded in primary and early auditory cortex (Fig. 3e,f), hinting at their role in the earliest stages of the auditory hierarchy (67). Predictions derived from the RNN models were encoded in the same channels, consistent with RNNs also capturing simple regularities, but also in higher-order regions (frontal and temporal associative cortex) (Fig. 3f). We thus propose the existence of dual temporal prediction processes relying on (1) low-level regularities in auditory areas and (2) contextual dependencies in higher-order regions. The brain extracts both low- and high-level regularities from its context to predict upcoming inputs. Notably, a large body of work implicates the dorsal auditory-motor pathway in temporal prediction, with delta- and beta-band motor activity conveying temporal predictions to auditory regions (25, 31–34). Our results are consistent with a sensorimotor contribution: temporal predictions were encoded in the sensorimotor cortex, in about a third of its speech-responsive channels (Fig. 3d,f and S4), stemming from both simple and contextually rich models (Fig. 3f), and with a predominantly temporal rather than content profile (Fig. 4b,c). The apparent modesty of this contribution relative to auditory regions should be interpreted with two caveats: intracerebral sampling is densest in temporal and perisylvian cortex and sparser over the dorsal pathway (Fig. S4), and our low-frequency encoding approach does not capture beta-band power dynamics, a central vehicle of motor-based temporal predictions (33). The role of the dorsal pathway in the contextual temporal predictions identified here, notably during active listening or speech-in-noise perception (68–70), remains an open question.

Crucially, although temporal and content predictions sometimes co-localize, reflecting the link between the duration and identity of units (71), they are largely encoded by distinct neural populations (Fig. 4c,d). This coupling also accounts for the variance shared by the two predictions (Fig. 4a), as part of the neural variance is attributable to either prediction. How content and temporal prediction may influence each other, notably given that content representations themselves carry temporal structure (72, 73), will be the subject of further inquiries. Together, these results suggest that the brain exploits different levels of temporal structure in addition to content information to continuously shape attention in time and better process low-level input (55, 74). Temporal prediction is therefore a distinct and central mechanism in speech processing, lying at the interface between high- and low-level processing and operating over linguistic and possibly acoustic units.

A few points merit further discussion. Now that we have established that a contextual temporal prediction is encoded, a critical remaining question is which unit of speech most needs to be temporally predicted for speech comprehension. Our results highlighted a relationship of (in)dependence between temporal and content prediction, a relationship that can only be defined for content-bearing units, the canonical example being the word. One could nonetheless imagine the same process applying to slower, prosody-related units, such as intonational units (75, 76). On the other hand, if temporal prediction is a sensory facilitation process, it may also operate over purely acoustic elements, as suggested by work on syllable-rate cortical oscillations and envelope tracking (29, 62). By design, our acoustic controls absorb any such contribution, which therefore remains untested here. Relatedly, since linguistic boundaries and acoustic edges are intrinsically coupled in natural speech, whether the predicted onsets are ultimately defined acoustically or linguistically cannot be fully resolved here. Another limitation concerns our stimulus, a single ten-minute excerpt narrated by one speaker: our findings should be generalized using longer and more varied listening material, spanning multiple texts, speakers and languages, given the cross-linguistic differences observed at the corpus level.

In demonstrating the richness and importance of temporal structure for natural speech processing, we hope to bridge the gap between temporal and content prediction and open new perspectives on the predictive coding of sensory sequences more broadly. The sequential structure of speech and the temporal dynamics of its processing have been extensively studied within the predictive coding framework (9, 73, 77). Yet the temporal structure of speech itself has rarely been treated as a source of information. By highlighting its contextual, dynamic and probabilistic nature, we advocate for repositioning the temporal structure within speech processing frameworks as a source of complex information in its own right. In this view, the brain takes advantage of every source of information available to predict upcoming input (64, 78).

## Materials and methods

### 1. Speech corpora

#### Training corpora

The French training corpus was derived from the *SynPaFlex-Corpus* dataset (79), which consists of 87 hours of French classical literature audiobooks, all narrated by the same female speaker. From this corpus, 50 hours of audio were arbitrarily selected and processed using the BAS web service (80) to obtain time-aligned transcriptions, including onset times for words, syllables, and phonemes. These onset annotations were then converted into three binary time series (one for each linguistic unit), indicating onset presence (1) or absence (0) at each time step. A sampling frequency of 50 Hz (temporal resolution of 20 ms) was chosen to capture the phonemic timescale (16, 81) while balancing temporal precision with computational efficiency.

The English training corpus was derived from the *LibriTTS* corpus (train-clean-100) (82), a multispeaker dataset comprising approximately 585 hours of English audiobooks. A 50-hour subset was arbitrarily selected and processed using the same procedure as for French, resulting in the same 3 binary time series.

#### Testing corpora

To validate and compare the performance of our models, we evaluated them on 100 speech segments, computing one AUC score per segment. It was essential to use material distinct from the training set, especially for French, where the training corpus included only a single narrator.

The French test corpus was compiled from publicly available audiobooks on Librivox. Ten audiobooks, each narrated by a different speaker (including both male and female voices), were selected. From each book, a 10-minute excerpt was extracted and processed using the same pipeline described for the training data. Each of these 10 excerpts was then subdivided into 10 shorter segments, resulting in a total of 100 test segments.

The English test corpus was built from portions of LibriTTS not used during training. As with the French data, 100 one-minute segments were extracted and processed using the same segmentation and preprocessing procedure.

### 2. Models of temporal prediction

#### Probability density function and Hazard rate

We begin our analysis with two probabilistic reference functions which serve as theoretical benchmarks for temporal prediction. These functions are derived from the structure of the temporal distributions of inter onset intervals (IOIs). They provide a principled baseline against which we later compare the performance of machine learning models. These functions are computed from the empirical distribution of IOI in a reference time series (the *training corpora*). The **probability density function (PDF)** is constructed by measuring the frequency of the different IOI values, yielding a discrete distribution over delays. For each prediction target, this distribution is then used to estimate the likelihood of observing an event at a given time step based on the time elapsed since the last event.

From the same distribution, we compute the **hazard rate (HR)** (41, 42, 49), which refines the PDF by incorporating the cumulative probability of event occurrence. The hazard rate expresses the instantaneous probability of an event occurring at a given time step, conditioned on the event *not* having occurred since the previous one. Formally, it is defined as:

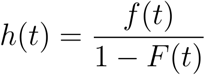

where *f*(*t*) is the probability density function (PDF) and *F*(*t*) is its cumulative distribution (CDF).

#### RNN-3D model

We designed a recurrent neural network to capture the temporal structure of natural continuous speech. The model *simultaneously* processes the onsets of *all* three linguistic units (words, syllables, and phonemes) as input (hence the name *RNN-3D*). It receives three binary time series, one per linguistic unit, indicating the presence (1) or absence (0) of an onset, sampled at 50 Hz. The model is trained to perform a multi-label prediction task, estimating the probability that an onset will occur for each linguistic unit at the next time step. Training was conducted on a 25-hour subset of the processed corpus of French (or English) speech, a duration at which performance plateaued.

RNN-3D was implemented using the PyTorch Python package (83). The model consists of a two-layer Gated Recurrent Unit (GRU) (84) network (input size = 3; hidden size = 20; dropout = 0.5) followed by a fully connected linear layer (output size = 3). Training was performed using batch learning (batch size = 500 time steps), with an Adam optimizer (learning rate = 0.001) for 100 epochs and binary cross entropy as the loss function. Model selection was based on the trade-off between training cost and test performance (AUC), rather than by loss convergence alone.

During the testing phase, to ensure that each prediction uses the same context length, all time series were segmented into overlapping sequences, using a sliding window of fixed length (sequence length = 500) and step size (step size = 1). Therefore, every prediction was made with only the previous 500 steps as context i.e. 10 seconds. This procedure was consistently used for all models included in the study.

#### RNN-1D model

The *RNN-1D* model shared the same architecture and training procedure as RNN-3D but was trained *separately* for each linguistic unit (phoneme, syllable, or word). Each model received a single binary onset time series as input, with both the input and output dimensions set to 1. All other parameters (GRU configuration, loss function, optimizer, and training setup) were identical to those used in RNN-3D.

### 3. Neuroimaging experiment

#### Participants

53 patients (25 females; mean age: 31 years; range: 17-54) with pharmacoresistant epilepsy took part in the study. All patients were French native speakers. They were implanted with depth electrodes for clinical purposes at the Hôpital de la Timone (Marseille, France) and at the Hôpital de l’Enfant Jésus, CHU de Québec - Université Laval (Canada, n=10). Neural recordings were performed 3 to 10 days post-implantation and always at least 4 hours after the last seizure. Patients were included in the study if their implantation map covered at least partially the Heschl’s gyrus (left or right). Neuropsychological assessments carried out before stereotactic electro-encephalographic (sEEG) recordings indicated that all patients had intact language functions and met the criteria for normal hearing. None of them had their epileptogenic zone including the auditory areas as identified by experienced epileptologists. Informed consent was obtained from all patients.

The study was approved by the Assistance Publique – Hôpitaux de Marseille (health data access portal registration number PADS 3ZSKYE) for the French data and by the Ethics Review Board of the CHU de Québec - Université Laval (2022-5890, Québec, QC, Canada) for the Canadian data. Recordings, interpretation and analysis of sEEG were performed following the French guidelines on stereoelectroencephalography (85) and recommendations on sEEG analysis (86).

#### Data acquisition

The sEEG signal was recorded using depth electrodes shafts of 0.8-mm diameter containing 6 to 18 electrode contacts (Dixi Medical or Alcis in France; AdTech in Canada). The contacts were 2 mm long and were spaced from each other by 1.5 mm (France) or 3-6 mm (Canada). The locations of the electrode implantations were determined solely on clinical grounds. Patients were recorded in their bedroom, using a digital 256-channel Natus amplifier (DeltaMed system), sampled at 512 Hz with 16-bits resolution, a hardware high-pass filter (cutoff = 0.16 Hz), and an antialiasing low-pass filter (cutoff = 340 Hz). The recording reference and ground were chosen by the clinical staff as two consecutive sEEG contacts on the same shaft both located in the white matter and/or at distance from any epileptic activity.

#### Preprocessing

Neural signals were corrected for DC shifts by using a high-pass filter with a 0.3-Hz cutoff frequency. Power line noise was attenuated using notch filters applied according to the geographical location of data acquisition: for recordings performed in France, a notch filter centered at 50 Hz was used; for recordings performed in Canada (n = 10), a notch filter was applied at 60 Hz. When necessary, additional notch filters were also applied at the respective harmonics of the power line frequency. In all cases, the notch bandwidth was defined as F₀/30 (default value in MNE-Python). To increase spatial sensitivity and reduce passive volume conduction from neighboring brain regions (86, 87), the signal was then offline referenced into a bipolar montage by subtracting activity recorded at each contact of interest from activity acquired at its closest neighbor site within the same electrode. This re-referencing process created virtual channels located at the midpoint of the original electrode locations. Finally, neural signals were band-pass filtered between 0.3 and 24 Hz.

To precisely localize the channels, a procedure similar to the one used in the iELVis toolbox was applied (88). First, we manually identified the location of each channel centroid on the post- implant CT scan using the Gardel software (89). Second, we performed volumetric segmentation and cortical reconstruction on the pre-implant MRI with the Freesurfer image analysis suite (90). In addition, segmentation of the pre-implant MRI with SPM12 provided the tissue probability maps (i.e. gray, white, and cerebrospinal fluid [CSF] probabilities) and the indexed-binary representations (i.e. either gray, white, CSF, bone, or soft tissues). Third, the post-implant CT scan was coregistered to the pre-implant MRI via a rigid affine transformation and the pre-implant MRI was registered to MNI152 space, via a linear and a non-linear transformation from SPM12 methods (91), through the FieldTrip toolbox (92). Each bipolar channel was then assigned to an anatomical region based on the Destrieux parcellation (93), grouped into eight regions of interest: auditory core, auditory belt, superior temporal gyrus (STG), superior temporal sulcus (STS), temporal associative cortex (TAC), insula, sensorimotor cortex (SMC) and frontal cortex (FC) (Fig. S4). All channels were included in the channel-wise analyses; channels falling outside these regions, including white-matter channels, were excluded from the regional analyses only.

#### Experimental design

Patients passively listened to approximately 10 minutes (577 s) of storytelling (La Sorcière de la rue Mouffetard; Gripari, 2004). The stimulus was presented via Sennheiser HD 25 headphones or loudspeakers, at a sound level of approximately 75 dBA, with a sampling rate of 44.1 kHz and 16-bit resolution.

#### Acoustic and onset Features

To ensure that the variance explained by our temporal prediction features could not be attributed to low-level acoustic or higher-level speech features (51, 94), we extracted a set of control features from the stimulus. Each was selected for its relevance to different acoustic dimensions of speech known to drive neural responses in the auditory cortex.

- Linguistic onsets: Onset times of words, syllables, and phonemes, estimated using the BAS web service (80). These onsets formed the binary time-series that fed the temporal-prediction models and were also entered as binary control regressors (Table 1).
- Spectrogram (8 frequency bands): To capture the spectro-temporal structure of the acoustic signal, we computed an auditory spectrogram using a gammatone filterbank, which approximates the frequency decomposition performed by the cochlea (95). The speech waveform was first decomposed into 128 narrow frequency bands spanning 20–5000 Hz, sampled at 1000 Hz (1-ms steps), and log-compressed (log(x + 1)). The 128 bands were then grouped into 8 broader frequency bands by summing across adjacent filters (16 filters per band). This yielded 8 regressors, each reflecting the amplitude fluctuations within one frequency band, providing a frequency-resolved representation richer than a single broadband envelope while still capturing the slow amplitude dynamics that approximate the syllabic timescale (16, 96).
- Spectrogram derivative (acoustic onsets, 8 frequency bands): For each of the 8 frequency bands, we additionally computed the half-wave-rectified first temporal derivative, defined as the positive part of the difference between successive time samples (94). This emphasizes transient increases in energy (acoustic onsets) within each band, which are strongly encoded in the auditory cortex and relate to the perception of acoustic edges and the syllabic rhythm (65).
- Intensity: This feature quantifies the acoustic energy of the signal averaged over short time frames and serves as a correlate of perceived loudness. It was computed in Praat (97) using a windowed root-mean-square (RMS) calculation, expressed in dB SPL with perceptual weighting for frequency sensitivity. Intensity closely tracks the broadband amplitude envelope (Fig. S2), and its slower fluctuations convey stress and prosodic prominence, signaling lexical and phrasal emphasis (98, 99).
- Pitch strength (also referred to as periodicity or voicing strength): The strength of the fundamental periodicity, extracted using Praat’s autocorrelation algorithm (100) as the per-frame strength of the selected pitch candidate, ranging from 0 (silent or unvoiced segments) to 1. It indexes the presence and salience of voicing over time, approximating the phonetic voicing contrast encoded in auditory cortex (101, 102)).
- Spectral flux: This measures the frame-to-frame variation in the spectral content of the signal, computed as the L2 norm (Euclidean distance) between successive normalized power spectra. It captures local spectral dynamics while being insensitive to global amplitude or phase, and is known to exhibit a bimodal spectral profile reflecting both syllabic and phonemic timescales (51).

#### Linguistic content Features

In addition to acoustic features, we included information-theoretic predictors quantifying the predictability of the speech input at the level of words and phonemes: surprisal (the unexpectedness of the observed unit in context) and entropy (the uncertainty of the prediction before the unit is observed) (4–6), resulting in 4 content features. Lexical and phonemic predictions were derived from CamemBERT (103), a transformer-based masked language model for

French (camembert-base). Words, phonemes, and their onsets were computed using BAS forced alignment (TextGrid MAU and ORT-MAU tiers), identically to temporal features.

For each word, predictions were obtained by masking the upcoming word and querying the model from its left context only, advancing the mask token by token. This procedure enforces strictly incremental (left-to-right) predictions, mimicking online comprehension despite the bidirectional architecture of the model. Word surprisal was computed as the mean negative log-probability across the upcoming word’s subword tokens:

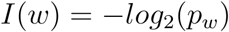

with *p_w_* the probability of word w. Word entropy as the Shannon entropy of the model’s predictive distribution at the word’s first token:

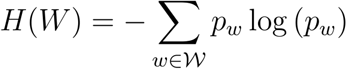

with *W* the distributions of possible words.

For phonemes, CamemBERT’s subword-token probabilities were mapped onto phonemes using a token-to-phoneme correspondence table: the probability of a phoneme was obtained by summing the probabilities of all tokens compatible with that phoneme at the current position, conditioned incrementally on the preceding phonemes of the word (a cohort scheme) and normalized over all phonemes admissible at that position. Phoneme surprisal was hence defined as the negative log-probability of the upcoming phoneme and phoneme entropy as the Shannon entropy of the normalized phoneme distribution, using the same formula as for words. Phonemes whose BAS segmentation diverged from the model’s token-based segmentation were marked as missing and set to zero in the corresponding regressors (i.e., treated as the absence of an event).

#### Temporal Features

The temporal features are the output of the different predictive models (RNN-1D, RNN-3D, PDF, and HR), for each linguistic unit i.e. the probability of occurrence of the different linguistic units (words, syllables, and phonemes) at each timestep.

In order to find the optimal representation of temporal prediction, we also computed features derived from these probabilities, namely the surprisal computed at each timestep, or continuous surprisal:

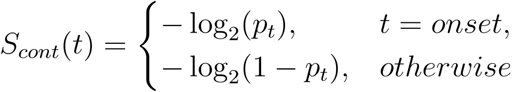

and the surprisal time-locked onto the corresponding unit onset, or discrete surprisal:

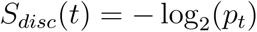

with *p_t_* the probability of onset occurrence at time t. Discrete surprisal is sparse (onset-aligned), whereas continuous surprisal is defined at every time step.

#### TRF analysis

We used the Temporal Response Function (TRF) framework to estimate how neural activity encodes different stimulus features, implemented as TRF regressors. All regressors were z-scored before being mapped to the neural data. All regressors and neural signals were resampled to 50 Hz for TRF estimation. TRF computations were performed using the sPyEEG library (50). A TRF is a linear model that, via temporal convolution, quantifies the relationship between continuous regressors and neural activity. For this analysis, the entire duration of the recordings was preserved, i.e., no artifacted epochs were excluded. When applied in a forward manner, the TRF describes the mapping of features onto the neural response (henceforth ‘encoding’; (104) as follows:

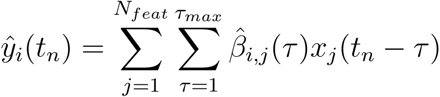

with *ŷ_i_*(*tn*) the estimated neural signal by the TRF model at time *t_n_*, *β_i,j_* the fitted coefficients of the TRF associated to the feature *i* and lag *j*.

The TRF was fitted using Ridge regression to avoid overfitting. For each patient, model performance was computed on held-out folds using 5-fold cross-validation across contiguous time segments and averaged across folds; for each channel, R²_cv was then taken at the regularization parameter (λ) maximizing this fold-averaged R². The same λ-selection procedure was applied within each permutation. Time lags from –0.2 s to 0.6 s were considered. The quality of the predicted neural response was assessed using the coefficient of determination (R²) between predicted and actual neural activity for each channel and model.

To partition the variance explained by temporal features, we built several TRF models. The control model included only the control features (acoustics, onsets and content features, see above). The global model (Fig. 3b) combined the control features with the three temporal features (word, syllable, phoneme). The overall contribution of temporal features was quantified as the difference in variance explained between the global and control models:

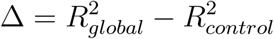

To estimate the unique contribution of each temporal feature, we compared the variance explained by the global model to that of a reduced model in which only the tested feature was excluded, while the other temporal features were preserved. For example, the unique contribution of the phoneme feature was:

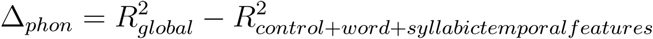

The same procedure was applied for the word and syllable features.

### 4. Statistics

#### All tests were two-sided

The performance of the onset prediction models (PDF, HR, RNN-1D, RNN-3D) was assessed using the area under the ROC curve (105). For each *model × unit* combination, 100 AUC scores were obtained from held-out excerpts. These scores were compared both against chance-level (AUC = 0.5) using Bonferroni-corrected Wilcoxon signed-rank tests, and across models or units using Friedman tests with Nemenyi-corrected post-hoc comparisons. Effect sizes are reported as rank-biserial correlations.

For the neural encoding analyses, the variance explained by the features of interest (ΔR² = R² of the full model minus that of the reduced model; see TRF analysis) was tested for each channel against a permutation-based null distribution. Null distributions were generated by permuting the tested regressors (n = 1000 permutations). Continuous regressors were circularly shifted in time by random offsets, preserving their temporal autocorrelation while abolishing their alignment with the neural signal. For onset-locked linguistic content regressors (surprisal and entropy), the values assigned to onsets were shuffled across onsets, preserving the temporal structure of events while abolishing the association between each event and its value. To control the family-wise error rate across channels, we used a maximum-statistic procedure within each participant: on every permutation, we retained the largest ΔR² across that participant’s channels, yielding a participant-level null distribution of maxima. A channel was considered significant when its observed ΔR² exceeded the 95th percentile of this max-distribution (family-wise error controlled at p < 0.05). The same procedure was used to identify channels significant for the control model alone (speech-responsive channels), except that it was performed on the R² scores rather than on ΔR² (no full-vs-reduced difference being defined for the control).

To compare **TRF encoding models** (PDF, HR, RNN-1D, RNN-3D), we used the proportion of significant channels per participant. An omnibus Friedman test (repeated measures across participants) was followed by pairwise Wilcoxon signed-rank tests between models ranked by decreasing mean (adjacent pairs only), Bonferroni-corrected by the number of comparisons. The same test was used for all model comparisons.

For the regional (atlas) breakdowns (model-family comparison (Fig. 3f); temporal-vs-content comparison (Fig. 4c)), we tested whether the proportion of channels exclusive to one family or condition differed between brain regions, using two-sided Fisher’s exact tests on every pair of regions, Bonferroni-corrected by the number of region pairs. For the model-family breakdown, the test contrasted RNN-only channels against the remaining channels (Combined + PDF/HR); for the temporal-vs-content breakdown, it contrasted Temporal-only against Content-only channels.

Finally, the proportion of significant channels per participant for the Temporal, Content, and Content+Temporal models was compared using pairwise Wilcoxon signed-rank tests across all three pairs, Bonferroni-corrected.

## Authors contribution

L.D. and P.H.G. designed research; L.D. performed research; L.D. and P.H.G. contributed new analytic tools; P.A. and A.T. collected the intracerebral data; L.D. analyzed data; D.S., B.M., and P.H.G. supervised research; L.D. wrote the first draft of the paper; B.M., and P.H.G. edited the paper; all authors read the paper.

## Acknowledgments

We thank all the members of D-CAP-INS for helpful discussions throughout this project. This work was co-funded by the European Union (ERC, SPEEDY, ERC-CoG-101043344), Fondation Pour l’Audition (FPA RD-2022-09), France 2030 (ILCB; ANR-16-CONV-0002) and the Excellence Initiative of Aix-Marseille University (A*MIDEX AMX-19-IET-004).

## Data, Materials, and Software Availability

All analysis code will be publicly available on GitHub upon publication (https://github.com/DCP-INS). The speech corpora used for model training and evaluation are publicly available (SynPaFlex, LibriTTS, LibriVox). The intracerebral EEG data cannot be publicly deposited owing to patient privacy regulations and the terms of the ethics approvals; deidentified data are available from the corresponding authors upon reasonable request.

## Supplementary Material

**Figure S1.**
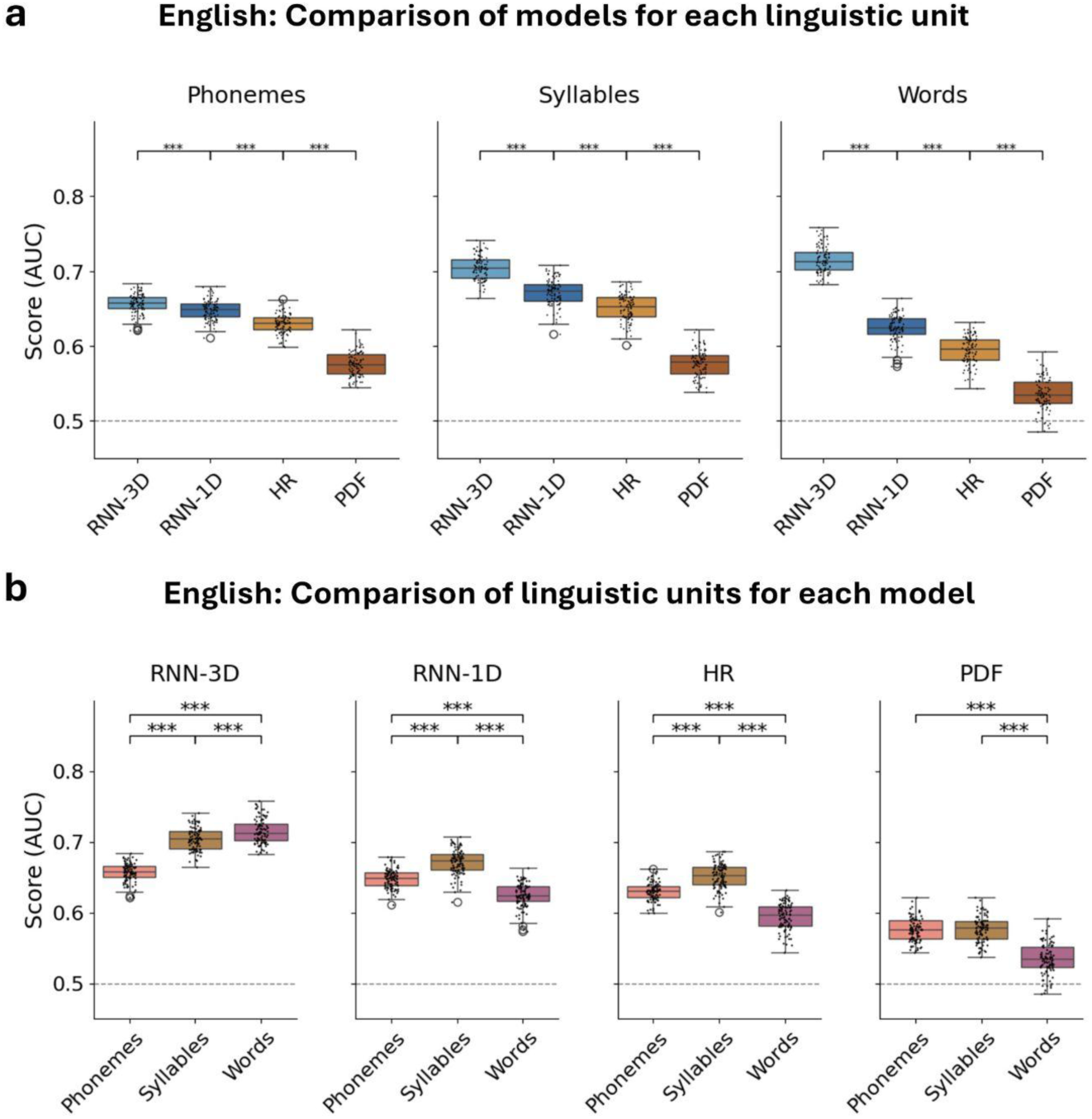
Comparison of model performance across linguistic units in English. Distribution of AUC scores computed on 100 one-minute speech segments. Models were evaluated based on their ability to temporally predict the onset of upcoming linguistic units (phonemes, syllables, or words). **(a)** AUC score comparison between models, for each linguistic unit. **(b)** AUC score comparison between linguistic units, for each model. Conventions as in Figure 2.

**Figure S2.**
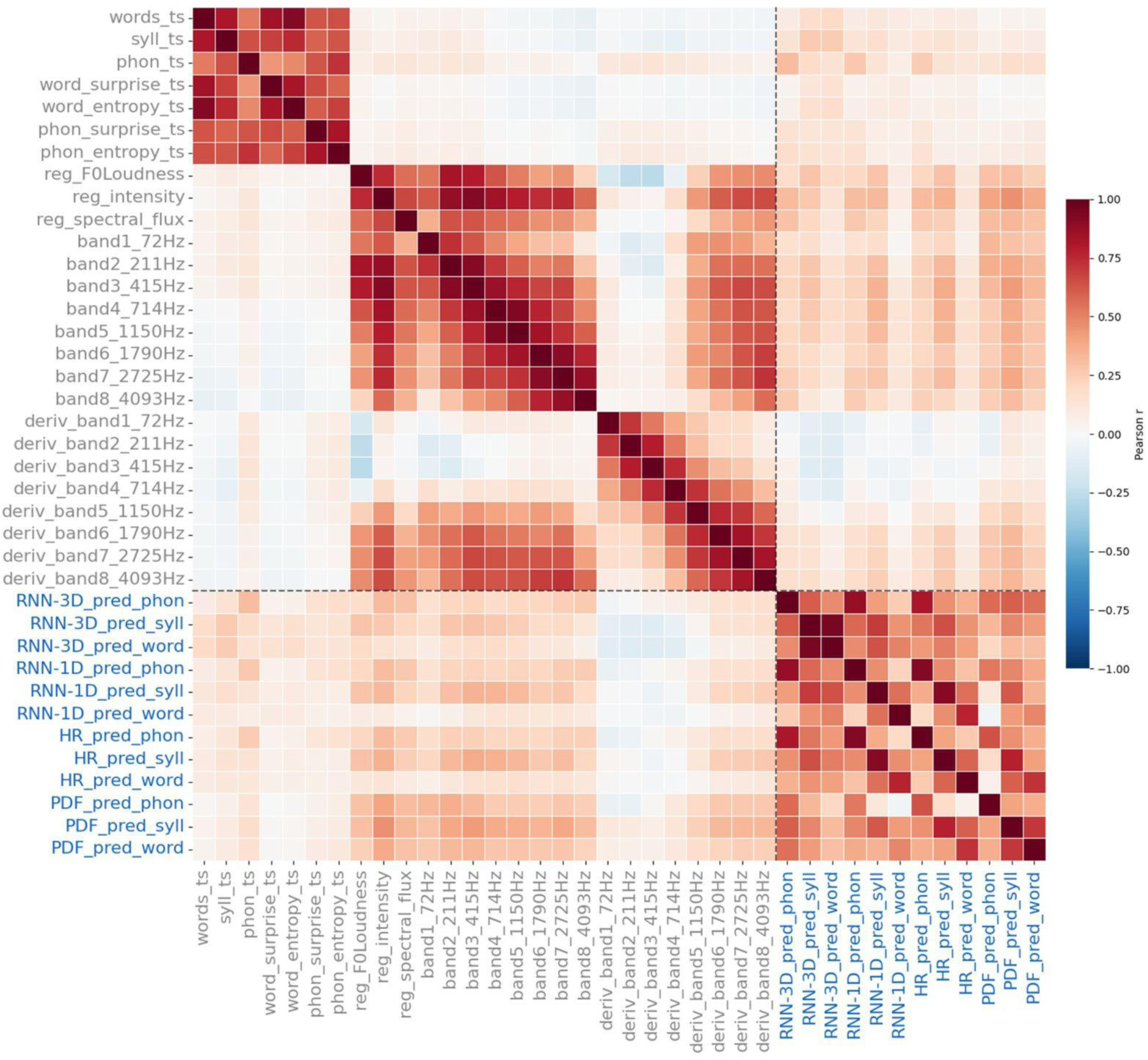
Correlation structure of the regressors used in the TRF analyses. Pearson correlation matrix (r) between all regressors, computed over all time steps. Regressors are grouped into control regressors (grey, n = 26) and regressors of interest (temporal prediction; blue, n = 12). The control set comprises the acoustic features (an 8-band spectrogram envelope and its 8-band derivative (acoustic edges), intensity, pitch strength (f0) and spectral flux), linguistic onset features (phoneme, syllable and word onsets), and linguistic content features (surprisal and entropy for words and phonemes). The regressors of interest are the onset-prediction outputs of the RNN-3D, RNN-1D, HR and PDF models for phonemes, syllables and words. The color scale runs from −1 (blue) to +1 (red).

**Figure S3.**
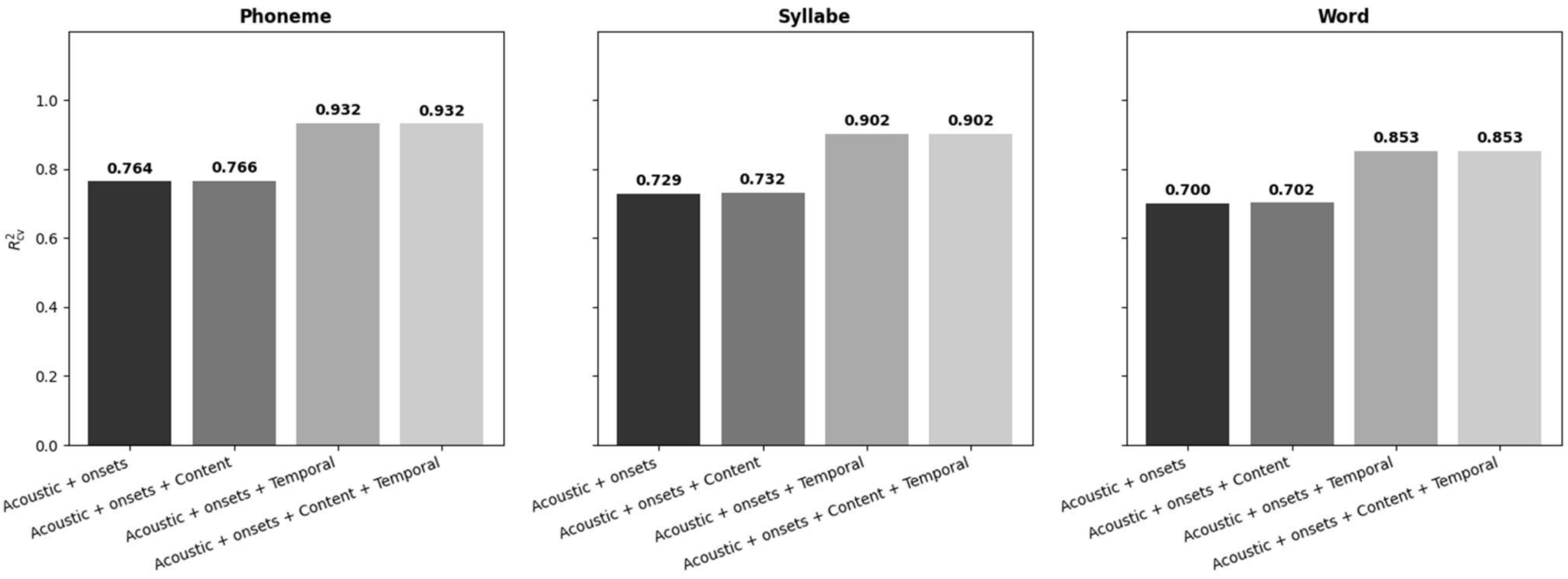
Variance of the RNN-1D regressor explained by the other regressor sets (TRF reconstruction). For each linguistic unit (Phoneme, Syllable, Word), a TRF was used to reconstruct the RNN-1D onset-prediction regressor from different (nested) sets of regressors, and the cross-validated reconstruction performance (R²_cv) is reported. Regressor sets: Acoustic (= Acoustics + Onsets; Table 1); Acoustic + Content; Acoustic + Temporal (Acoustics + Onsets + the other temporal-prediction regressors RNN-3D, PDF and HR, i.e. excluding RNN-1D); and Acoustic + Content + Temporal. No permutation tests were performed for this figure.

**Figure S4.**
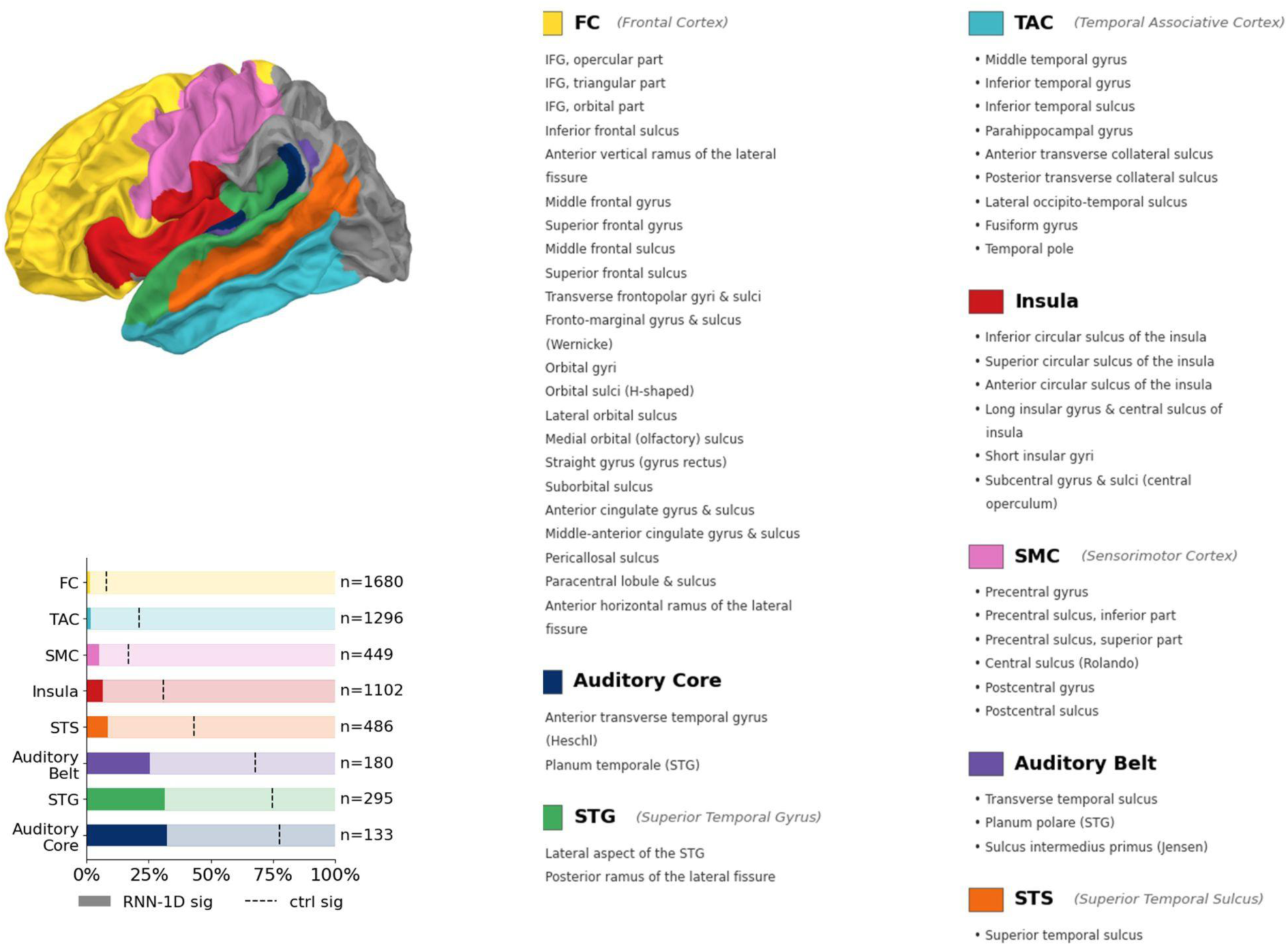
Anatomical parcellation and regional distribution of significant channels (related to Figs. 3 and 4). **Left.** Reconstructed cortical surface showing the anatomical regions used to group channels in Figures 3 and 4. **Right.** For each region, the proportion of channels significant for the RNN-1D model (filled bar) out of all channels located in that region (pale bar = 100% of the region’s channels); the dashed vertical line marks the proportion of channels significant for speech-responsive channels (Acoustics + Onsets + Content; Table 1) only. Regions are ordered by increasing RNN-1D proportion; n indicates the total number of channels in the region. Channel significance was assessed with an FWER-corrected permutation test (max statistic, n = 1000, p ≤ 0.05). Region abbreviations: Auditory Core, Auditory Belt, STG (superior temporal gyrus), STS (superior temporal sulcus), TAC (temporal associative cortex), Insula, SMC (sensorimotor cortex), FC (frontal cortex); channels falling outside these regions were excluded. The parcellation is based on the Destrieux atlas (FreeSurfer aparc.a2009s; Destrieux et al., 2010), projected onto the fsaverage surface.

**Figure S5.**
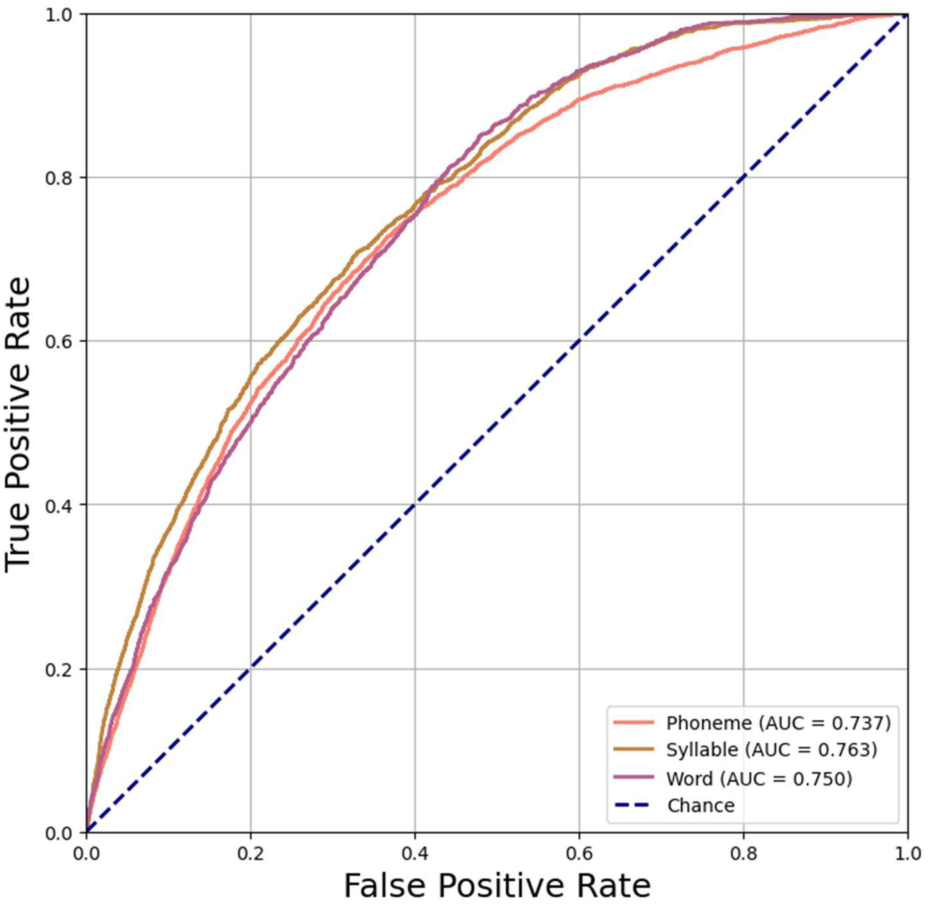
ROC curves illustrating the performance of the RNN-3D model on the audiobook used in the intracerebral EEG recording experiment. The model was trained on 25h of French audiobook corpus data on all linguistic units, and tested on its ability to predict phoneme, syllable, and word onsets in the 10-minute French audiobook. Each curve shows the true positive rate as a function of the false positive rate for a given linguistic unit, with the corresponding area under the curve (AUC) values reported in the legend. The RNN-3D model is the best-performing onset predictor at the corpus level (Fig. 2).

**Figure S6.**
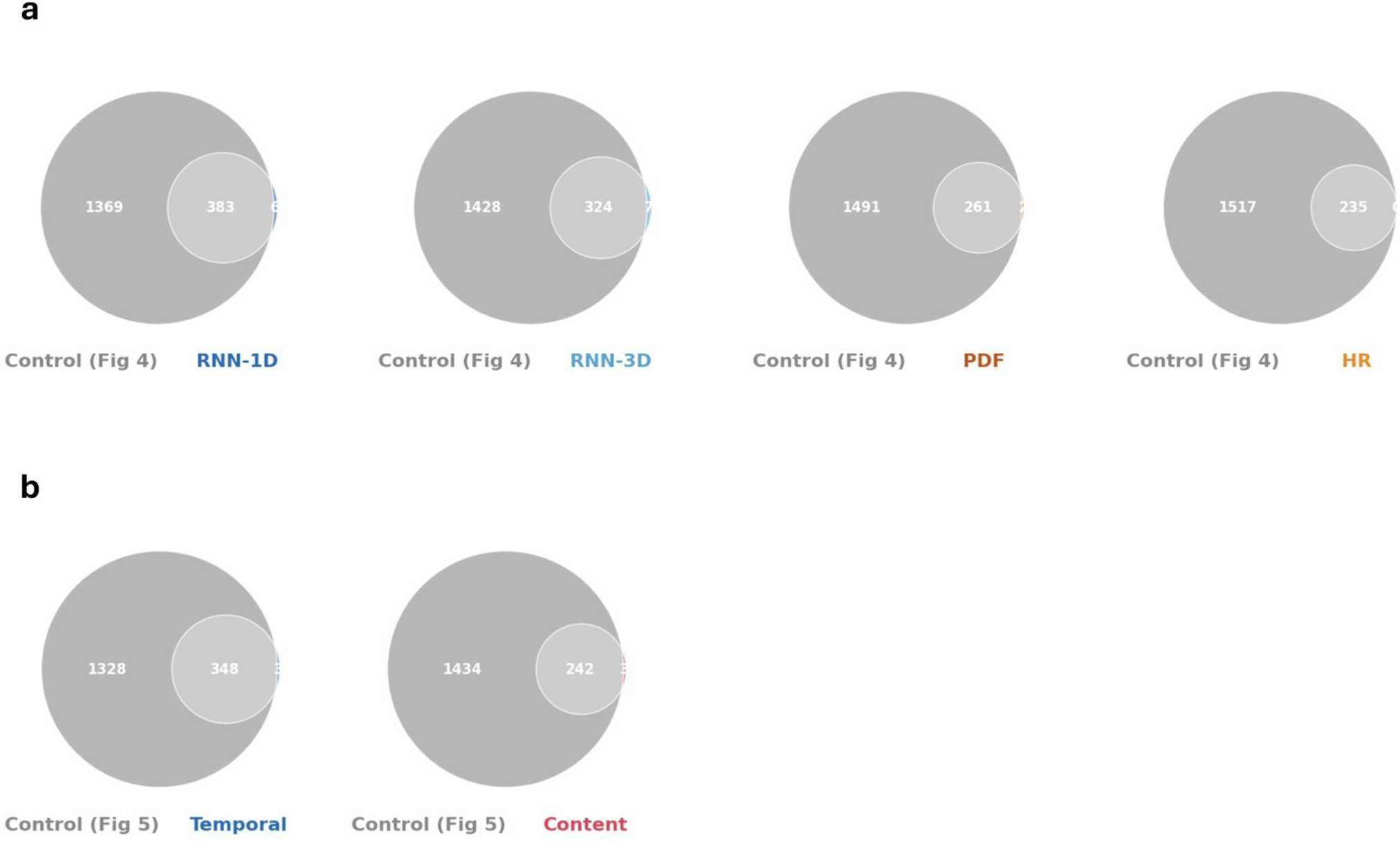
Significant model channels are a subset of the channels significant for their respective control model (related to Figs. 3 and 4). **(a)** Venn diagrams comparing, across all participants, the set of channels significant for the control model in Fig. 3 (Acoustics + Onsets + Content; Table 1; speech-responsive channels) with the set of channels significant for temporal-prediction models (RNN-1D, RNN-3D, PDF, HR). **(b)** Venn diagrams comparing, across all participants, the set of channels significant for the control model in Fig. 4 (Acoustics + Onsets; Table 2; control-significant channels) with the set of channels significant for the Temporal and Content models in Fig. 4. **(a and b)** Channel significance was assessed with an FWER-corrected permutation test (max statistic, n = 1000, p ≤ 0.05). Significance for the non-control models reflects the unique variance of the added regressors (beyond control), hence the subset relationship.

**Figure S7.**
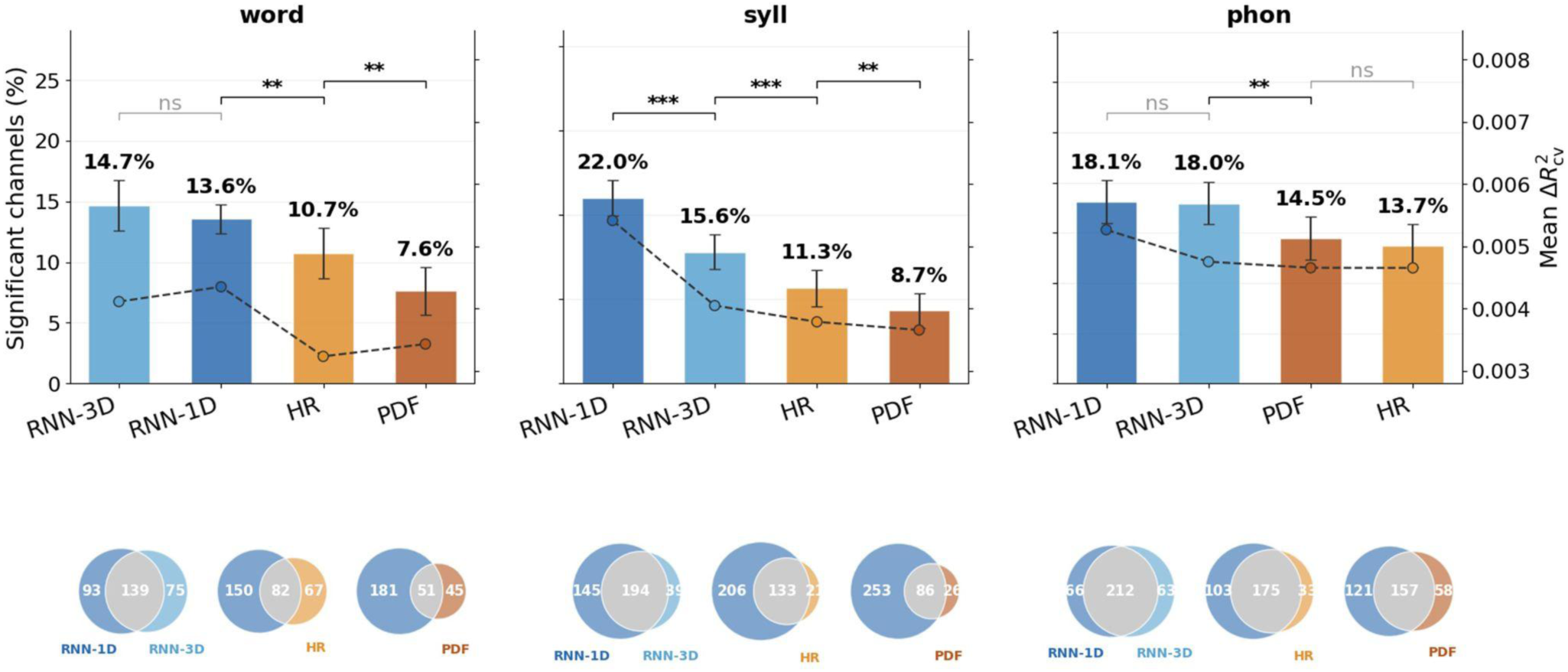
Comparison of model performance for each linguistic unit separately (related to Fig. 3b). The same analysis as in Figure 3b was performed independently for word, syllable (syll), and phoneme (phon) onset predictions (dataset: 7,698 channels, 53 participants). **Top.** Percentage of speech-responsive channels significant for each model (mean ± SEM across participants). Bars are ordered by decreasing mean within each unit. The mean unique variance (ΔR²_cv, right axis, dashed line) is shown for reference. **Bottom.** Venn diagrams showing the overlap in significant channels between RNN-1D and each of the three other models (RNN-3D, HR, PDF), pooled across participants. Conventions as in Figure 3.

**Figure S8.**
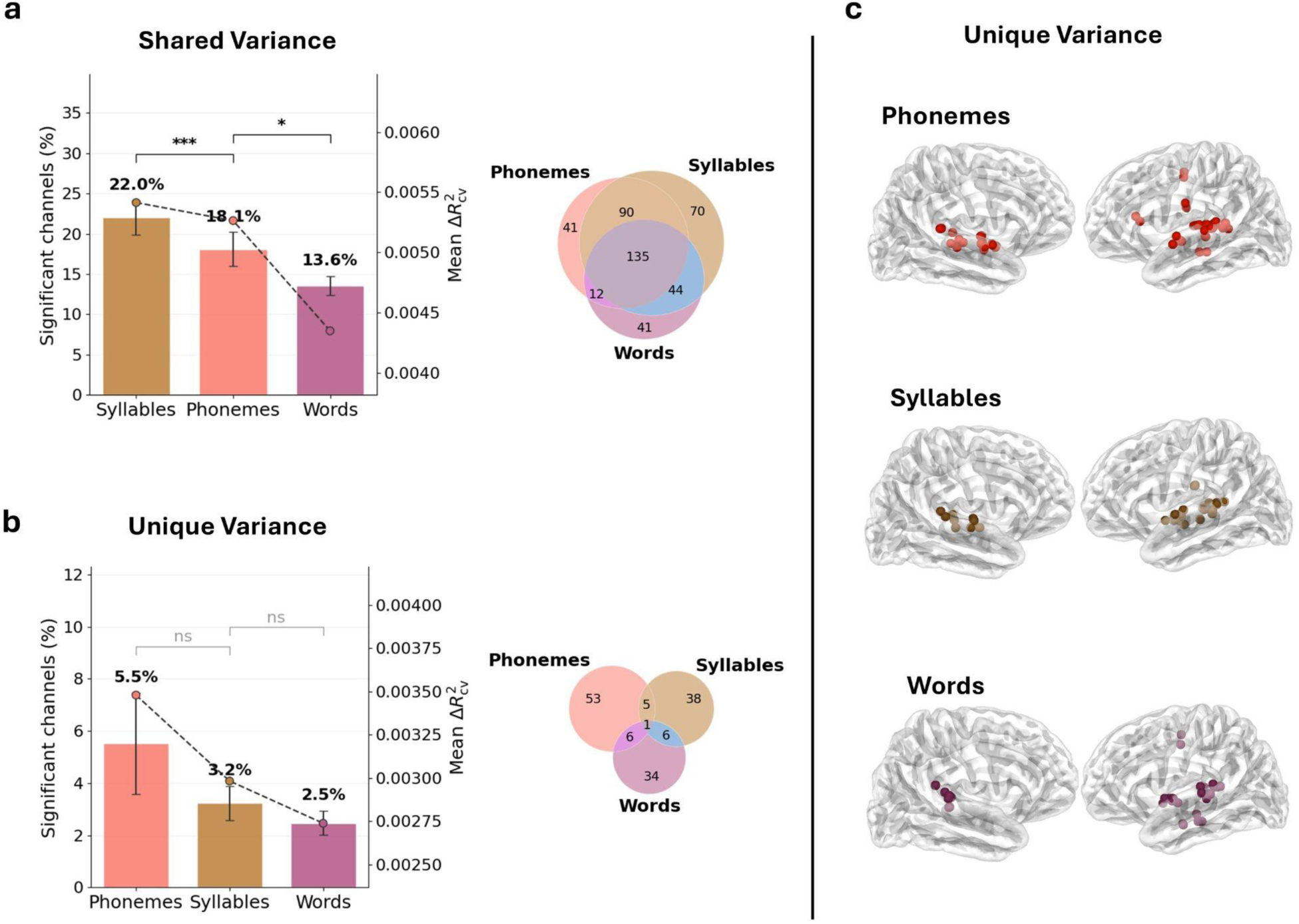
Shared and unique variance of the RNN-1D temporal prediction across linguistic units (related to Fig. 3b and Fig. S73). RNN-1D model only, comparing its three linguistic units (word, syllable [syll], phoneme [phon]), ordered by decreasing mean. In panels (a) and (b), bars show the percentage of speech-responsive channels significant for each unit (mean ± SEM across participants). The dashed line (right axis) shows the mean ΔR²_cv. Venn diagrams show the overlap in significant channels across the three units (pooled across participants). Across-unit comparisons: Friedman test followed by pairwise Wilcoxon signed-rank tests on adjacent units, Bonferroni-corrected. Conventions as in Figure 3. **(a)** Shared variance: each unit is evaluated in a separate model (control [Acoustics + Onsets + Content; Table 1] + that unit), its contribution isolated by permuting the unit; this gain includes the variance the unit shares with the other units. **(b) Left.** Unique variance: each unit is evaluated within the full model (control + all three units), isolated by permuting only the tested unit, leaving the variance unique to it. **(c)** Anatomical distribution of the unique-variance channels (phonemes, top; syllables, middle; words, bottom), projected onto a standard cortical surface (left and right hemispheres).

**Figure S9.**
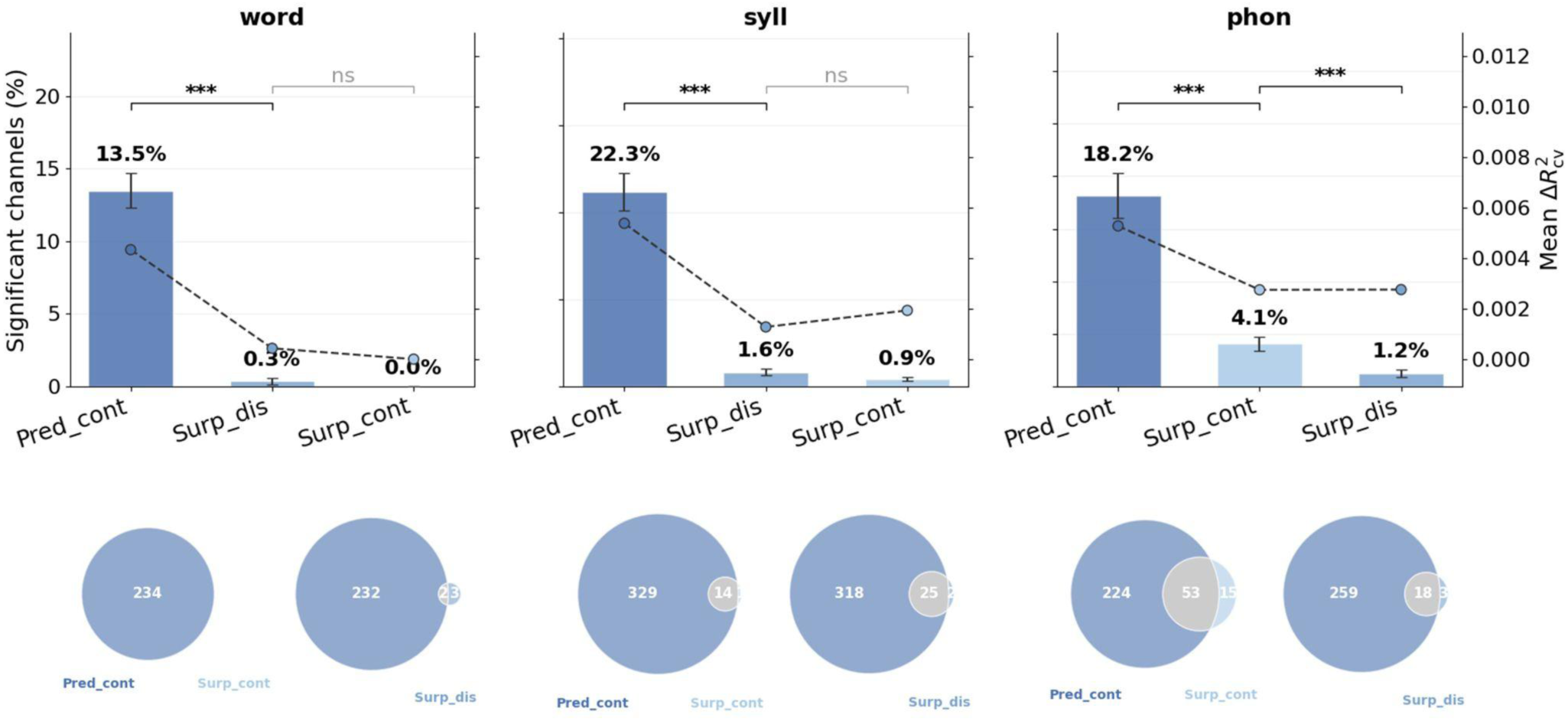
Comparison of predictive metrics for each linguistic unit (related to Fig. 3). This analysis verifies that the continuous onset-prediction signal (Pred_cont) is the most appropriate metric for quantifying the models’ temporal predictions used in the neural analyses of Figure 3. For each unit (word, syllable (syll), phoneme (phon)), the continuous prediction is compared with two surprisal-based metrics derived from the same model output: Surp_dis (discrete surprisal) and Surp_cont (continuous surprisal). Surprisal was computed as for the content metrics (Methods); in the continuous case it is defined as −log(pred) at onset time steps and −log(1−pred) at non-onset time steps (i.e. how surprising it is for an event not to occur). **Top.** Percentage of speech-responsive channels per participant significant for each metric (mean ± SEM across participants). Bars are ordered by decreasing mean within each unit. The mean unique variance (ΔR²_cv, right axis, dashed line) is shown for reference. Across-metric comparisons: Friedman test, then pairwise Wilcoxon on adjacent metrics, Bonferroni-corrected. **Bottom.** Venn diagrams of the overlap in significant channels between Pred_cont and each surprisal metric, pooled across participants. The continuous prediction captures substantially more significant channels than either surprisal metric across all units. Conventions as in Figure 3.

